# Suppression of p16 induces mTORC1-mediated nucleotide metabolic reprogramming

**DOI:** 10.1101/393876

**Authors:** Raquel Buj, Chi-Wei Chen, Erika S. Dahl, Kelly E. Leon, Ross Kuskovsky, Natella Maglakelidze, Maithili Navaratnarajah, Gao Zhang, Mary T. Doan, Helen Jiang, Michael Zaleski, Lydia Kutzler, Holly Lacko, Yiling Lu, Gordan B. Mills, Raghavendra Gowda, Gavin P. Robertson, Joshua I. Warrick, Meenhard Herlyn, Yuka Imamura, Scot R. Kimball, David J. DeGraff, Nathaniel W. Snyder, Katherine M. Aird

## Abstract

Reprogrammed metabolism and cell cycle dysregulation are two cancer hallmarks. p16 is a cell cycle inhibitor and tumor suppressor that is upregulated during oncogene-induced senescence (OIS). Loss of p16 allows for uninhibited cell cycle progression, bypass of OIS, and tumorigenesis. Whether p16 loss affects pro-tumorigenic metabolism is unclear. We report that suppression of p16 plays a central role in reprogramming metabolism by increasing nucleotide synthesis. This occurred via activation of mTORC1 signaling, which directly mediated increased translation of the mRNA encoding ribose-5-phosphate isomerase A (*RPIA*), a pentose phosphate pathway enzyme. p16 loss correlated with activation of the mTORC1-RPIA axis in multiple cancer types. Suppression of RPIA inhibited proliferation only in p16-low cells by inducing senescence both *in vitro* and *in vivo*. These data reveal the molecular basis whereby p16 loss modulates pro-tumorigenic metabolism through mTORC1-mediated upregulation of nucleotide synthesis and reveals a metabolic vulnerability of p16-null cancer cells.

**Highlights:**

- mTORC1 is activated by p16 knockdown to increase nucleotide synthesis and bypass senescence
- mTORC1 directly increases translation RPIA to increase ribose-5-phosphate
- Activation of mTORC1 pathway downstream of p16 suppression is independent of RB
- RPIA suppression induces senescence only in cells and tumors with low p16

## Introduction

Metabolic reprogramming is a hallmark of cancer (Hanahan and Weinberg, 2011; Pavlova and Thompson, 2016). Transformed and tumorigenic cells require increased deoxyribonucleotide synthesis to fuel the genome replication that sustains their unregulated cell cycle and proliferation. Therefore, it is likely that the cell cycle and nucleotide metabolism are linked. The cell cycle inhibitor p16 is a critical tumor suppressor that is lost as an early event in many human cancers (Belinsky et al., 1998; Chin, 2003; Hruban et al., 2000; Nuovo et al., 1999). Indeed, expression of p16 is low or null in approximately half of all human cancers (Li et al., 2011). This mostly occurs through homozygous deletion or DNA methylation and loss of heterozygosity (LOH) (Merlo et al., 1995; Ortega et al., 2002). The Cancer Genome Atlas (TCGA) shows 24% of melanomas and 28% of pancreatic cancers harbor homozygous deletions in cyclin dependent kinase inhibitor (*CDKN2A,* encoding for p16) (Cerami et al., 2012; Gao et al., 2013; Shain et al., 2018; Shain et al., 2015). In other cancers, such as colorectal cancers, *CDKN2A* is often silenced by promoter hypermethylation (12-51% of cases) (Herman et al., 1995; Shima et al., 2011). While loss of p16 is known to play a role in deregulating the cell cycle, whether loss of p16 expression affects nucleotide metabolism is unknown.

Both increased expression of p16 (Serrano et al., 1997) and decreased levels of dNTPs (Aird et al., 2013; Mannava et al., 2013) are characteristics of cellular senescence, a stable cell cycle arrest (Aird and Zhang, 2014, 2015; Dorr et al., 2013; Hernandez-Segura et al., 2018; Wiley and Campisi, 2016). Activation of oncogenes such a BRAF^V600E^ induces senescence to suppress transformation and tumorigenesis (termed oncogene-induced senescence, OIS) (Perez-Mancera et al., 2014; Yaswen and Campisi, 2007). Therefore, OIS is considered an important tumor suppressor mechanism *in vivo* (Braig et al., 2005; Michaloglou et al., 2005). Moreover, increased dNTPs or loss of p16 bypasses OIS to allow for transformation and tumorigenesis (Aird et al., 2015; Aird et al., 2013; Damsky et al., 2015; Dankort et al., 2007; Goel et al., 2009; Haferkamp et al., 2008; Sarkisian et al., 2007). Thus, we reasoned that these two processes may be interconnected.

Here we used senescence as a model to study the link between p16 and nucleotide metabolism. We found that depletion of p16 increases deoxyribonucleotide synthesis to bypass senescence induced by multiple stimuli, including dNTP depletion and BRAF^V600E^ expression. Mechanistically we determined that loss of p16 increases mTORC1-mediated translation of the pentose phosphate pathway (PPP) enzyme ribose-5-phosphae isomerase (RPIA) to upregulate production of ribose-5-phosphate and nucleotides. Underscoring the importance of this pathway in human cancers, mTORC1 activation correlates with decreased p16 expression and worse prognosis in multiple cancer types. Additionally, cancer cells with low p16 expression are more sensitive to the mTORC1 inhibitor temsirolimus and rely upon RPIA protein expression for proliferation both *in vitro* and *in vivo*. These data demonstrate that loss of p16 increases deoxyribonucleotide synthesis through upregulation of mTORC1 activity.

## Results

### p16 knockdown enhances nucleotide synthesis to bypass senescence

Loss of p16 is an early event in the progression from senescent benign lesions to cancer (Bennecke et al., 2010; Bennett, 2016; Caldwell et al., 2012; Kriegl et al., 2011; Michaloglou et al., 2005; Shain et al., 2015). We and others have demonstrated that increased dNTPs also bypasses senescence (Aird et al., 2013; Mannava et al., 2013). However, it is unknown whether these two events are linked. To determine whether p16 loss affects nucleotide synthesis, we took advantage of our previously-published model of dNTP-depletion induced-senescence by knocking down RRM2 (Aird et al., 2013). We have previously extensively validated this short hairpin RNA (shRNA) (Aird et al., 2013). Knockdown of p16 in shRRM2 cells (**Fig. 1A and S1A**) suppressed senescence markers including BrdU incorporation, colony forming ability, and senescence-associated beta-galactosidase (SA-β-Gal) activity (**Fig. 1B-E**). Data using a second independent hairpin targeting p16 and overexpression of p16 cDNA (**Fig. S1B-C**) demonstrate these results are specific for p16 (**Fig. S1D-K**). In order to further verify this observation in a pathologically-relevant model, we used BRAF^V600E^-induced senescence and knocked down p16 (**Fig. 1F**). Similar to our dNTP-depletion-induced senescence model, knockdown of p16 bypassed BRAF^V600E^-induced senescence (**Fig. 1G-J**). To determine whether knockdown of p16 altered dNDP/dNTP levels, we next determined the relative abundance of these deoxyribonucleotides by LC-HRMS. Knockdown of p16 in both senescence models significantly increased dNDPs/dNTPs even above control levels in some nucleotides (**Fig. 1K-L**). Note that dGDP/dGTP was not quantified due to spectral overlap with the highly abundant dADP/dATP. Interestingly, we observed an increase in RRM2B in shRRM2/shp16 cells (**Fig. S1L-M**), which is likely how these cells are able to reduce NDPs/NTPs to dNDPs/dNTPs. Gene Set Enrichment Analysis (GSEA) of two publicly available data sets (Kabbarah et al., 2010; Talantov et al., 2005) showed a significant enrichment in terms associated with nucleotide synthesis in melanoma when compared with human nevi (**Table S1**), which are considered senescent (Michaloglou et al., 2005), suggesting that nucleotide metabolism is relevant for senescence bypass *in vivo*. Excitingly, further metabolite analysis demonstrated that nucleotides were also significantly increased upon p16 knockdown in these models (**Fig. 1M-N and Fig. S1N**), suggesting that the increase in deoxyribonucleotides is not simply due to increased RRM2B or the proportion of cells in S-phase. Together, these data indicate that p16 depletion increases both nucleotide and deoxyribonucleotide synthesis to bypass senescence.

**Figure 1.**
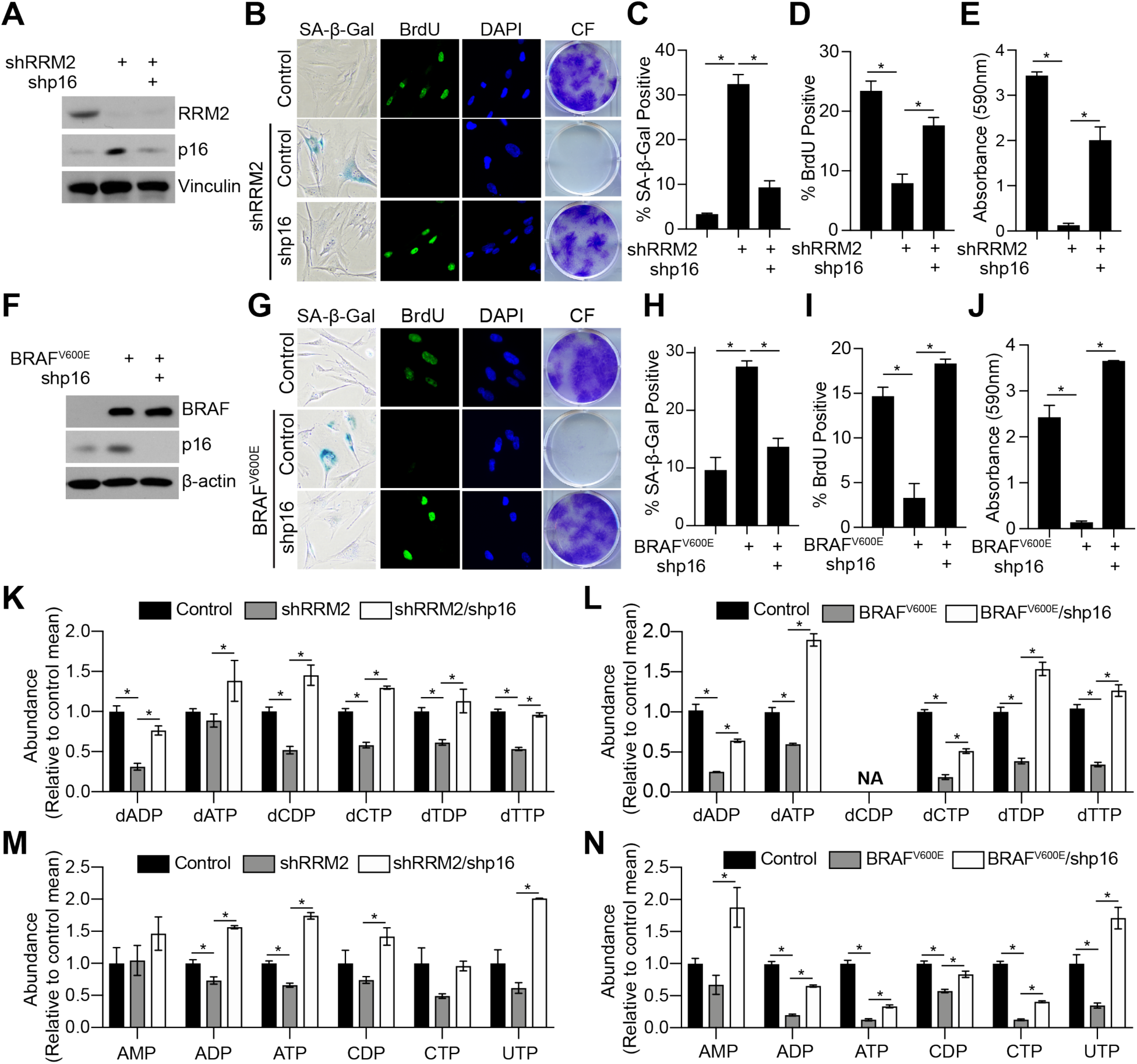
Suppression of p16 increases nucleotide synthesis to bypass senescence. **(A-E)** Normal diploid IMR90 cells were infected with lentivirus expressing short hairpin RNAs (shRNAs) targeting RRM2 (shRRM2) alone or in combination with an shRNA targeting p16 (shp16). Empty vector was used as a control. **(A)** Immunoblot analysis of indicated proteins. One of 5 experiments is shown. **(B)** SA-β-Gal activity, BrdU incorporation, and colony formation. One of 5 experiments is shown. **(C)** Quantification of SA-β-Gal activity in (B). n=3/group, one of 5 experiments is shown. Data represent mean ± SD. *p<0.001 **(D)** Quantification of BrdU incorporation in (B). n=3/group, one of 5 experiments is shown. Data represent mean ± SEM. *p<0.001 **(E)** Quantification of colony formation in (B). n=3/group, one of 5 experiments is shown. Data represent mean ± SEM. *p<0.001 **(F-J)** Normal diploid IMR90 cells were infected with retrovirus expressing BRAF^V600E^ alone or in combination with an shRNA targeting p16 (shp16). Empty vector was used as a control. **(F)** Immunoblot analysis of indicated proteins. One of 3 experiments is shown. **(G)** SA-β-Gal activity, BrdU incorporation, and colony formation. One of 5 experiments is shown. **(H)** Q uantification of SA-β-Gal activity in (G). n=3/group, one of 5 experiments is shown. Data represent mean ± SD. *p<0.001 **(I)** Quantification of BrdU incorporation in (G). n=3/group, one of 5 experiments is shown. Data represent mean ± SEM. *p<0.002 **(J)** Quantification of colony formation in (G). n=3/group, one of 5 experiments is shown. Data represent mean ± SEM. *p<0.001 **(K-N)** Nucleotide analysis was performed by LC-HRMS in the indicated groups. n>3/group, one of at least 2 experiments is shown. Data represent mean ± SEM. *p<0.05 NA= not available

### p16 knockdown activates mTORC1 to bypass senescence and increase nucleotide synthesis

We next aimed to determine the underlying mechanism of nucleotide synthesis upon p16 knockdown. p16 inhibits E2F-mediated transcription in part through regulating the retinoblastoma protein (RB)-E2F interaction (Sherr, 2001). Thus, we performed RNA-Seq (GSE133660). Interestingly, while we did not observe terms related to purine and pyrimidine synthesis, GSEA showed an enrichment in the mTORC1 signaling pathway in shRRM2/shp16 when compared with shRRM2 alone **(Table S2)**. The increase in mTOR signaling was confirmed by Reverse Phase Protein Array analysis **(Table S3)**. Underscoring the pathological relevance of our findings, the mTORC1 signaling pathway was also enriched in melanoma when compared with nevi in both Talantov and Kabbarah data sets **(Table S1)**. Previous studies have demonstrated that mTORC1 increases both purine and pyrimidine synthesis (Ben-Sahra et al., 2013; Ben-Sahra et al., 2016), suggesting that this may be the mechanism by which loss of p16 increases nucleotides in senescence bypass. We confirmed the activation of mTORC1 signaling in shRRM2/shp16 and BRAF^V600E^/shp16 by assessing the increased phosphorylation of S6K and 4E-BP1 (**Fig. 2A-B**) as well as by mTORC localization at the lysosomal membrane **(Fig. S2A-B).** Underscoring the role of mTORC1 promoting dNTP synthesis downstream of p16 loss, inhibition of mTORC1 with temsirolimus (**Fig. 2A-B**) significantly decreased both nucleotides and deoxyribonucleotides in shRRM2/shp16 and BRAF^V600E^/shp16 (**Fig. 2C-D and Table S4**). Consistent with the notion that the nucleotide synthesis downstream of mTORC1 is critical for cells to bypass senescence, mTORC1 inhibition with temsirolimus inhibited this phenotype (**Fig. 2E-L**). Temsirolimus dose had no effect on parental cells (**Fig. S2C**), suggesting that only cells with high mTORC1 activity are sensitive to the drug. Together, these data demonstrate that activation of mTORC1 downstream p16 loss drives the observed increase in dNDPs/dNTPs.

**Figure 2.**
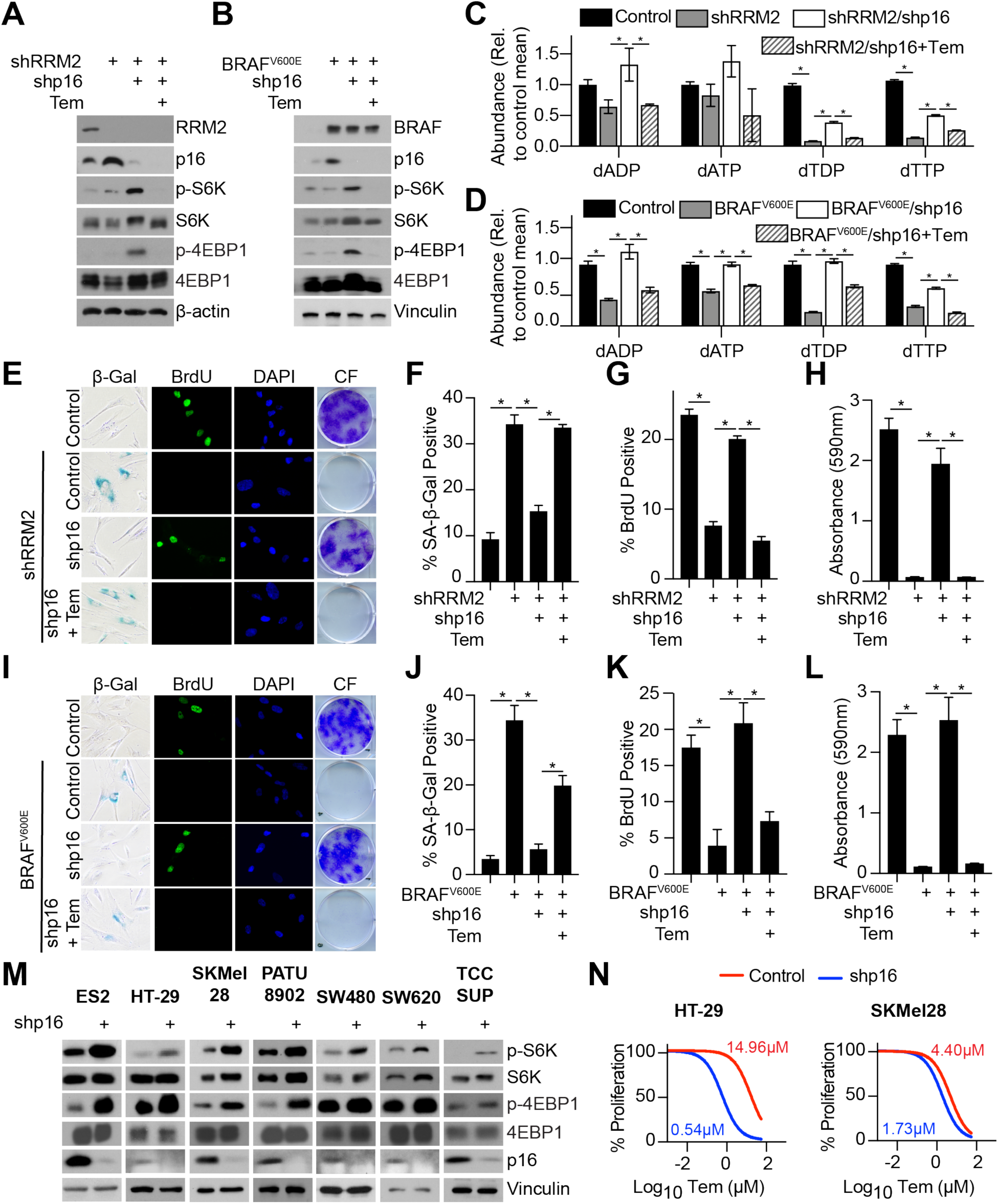
Suppression of p16 activates mTORC1 to increase nucleotide synthesis. **(A)** Normal diploid IMR90 cells were infected with lentivirus expressing short hairpin RNA (shRNAs) targeting RRM2 (shRRM2) alone or in combination with an shRNA targeting p16 (shp16). Empty vector was used as a control. Temsirolimus (Tem; 0.5nM) was added 4 days after starting selection (**Fig. S1A**). Immunoblot analysis of the indicated proteins. One of 3 experiments is shown. **(B)** Normal diploid IMR90 cells were infected with retrovirus expressing BRAF^V600E^ alone or in combination with an shRNA targeting p16 (shp16). Empty vector was used as a control. Temsirolimus (Tem; 0.5nM) was added 4 days after starting selection (**Fig. S1A**). Immunoblot analysis of the indicated proteins. One of 3 experiments is shown. **(C-D)** dNDP/dNTP analysis was performed by LC-HRMS in the indicated groups. n>3/group, one of at least 3 experiments is shown. Data represent mean ± SEM. *p<0.05 **(E)** Same as (A). SA-β-Gal activity, BrdU incorporation, and colony formation. One of 3 experiments is shown. **(F)** Quantification of SA-β-Gal activity in (E). n=3/group, one of 3 experiments is shown. Data represent mean ± SEM. *p<0.001 **(G)** Quantification of BrdU incorporation in (E). n=3/group, one of 3 experiments is shown. Data represent mean ± SEM. *p<0.001 **(H)** Quantification of colony formation in (E). n=3/group, one of 3 experiments is shown. Data represent mean ± SEM. *p<0.001 **(I)** Same as (B). SA-β-Gal activity, BrdU incorporation, and colony formation. One of 3 experiments is shown. **(J)** Quantification of SA-β-Gal activity in (I). n=3/group, one of 3 experiments is shown. Data represent mean ± SEM. *p<0.02 **(K)** Quantification of BrdU incorporation in (I). n=3/group, one of 3 experiments is shown. Data represent mean ± SEM. *p<0.001 **(L)** Quantification of colony formation in (I). n=3/group, one of 3 experiments is shown. Data represent mean ± SEM. *p<0.001 **(M)** The indicated cancer cell lines with wildtype p16 expression were infected with a short hairpin targeting p16. Cells were serum starved for 16h after which they were incubated with 10% FBS for 30 min. Immunoblot analysis of the indicated proteins. One of at least 2 experiments is shown. **(N)** Same as (M) but cells were treated with a dose-course of temsirolimus under 0.5% FBS conditions. n=3/group, one of 2 experiments is shown. Data represent non-linear fit of transformed data. IC_50_ for each condition is indicated.

mTORC1 is a master regulator of translation and mRNA metabolism (Ma and Blenis, 2009; Nandagopal and Roux, 2015). Interestingly, in TCGA patient samples, increased expression of leading-edge genes associated with “translation” GSEA term (**Table S2 and S5**) significantly co-occurred with alterations in *CDKN2A*, and this signature was associated with worse overall survival (**Fig. S2D-E**), highlighting the clinicopathological implications of this pathway. Because p16 expression is lost in multiple human cancers, we aimed to determine whether loss of p16 correlates with mTORC1 activation in tumor cells. We knocked down p16 in seven tumor cell lines from multiple cancer types with wildtype p16 expression. According to TCGA, all but TCCSUP also contain wildtype *RB1*. We observed increased phosphorylation of S6K and 4E-BP1 in all cell lines tested (**Fig. 2M**). Consistently, cancer cells with p16 knockdown were more sensitive to inhibition of mTORC1 (**Fig. 2N and Fig. S2F**), although the degree of sensitivity varied between cell lines likely due to genetic heterogeneity of the cell lines tested. Interestingly, TCCSUP had only a 2-fold decrease in IC_50_, which may be related to its *RB1* mutation. Finally, analysis of data from the Dependency Map (depmap.org) also indicates that cancer cells with low *CDKN2A* copy number are more sensitive to temsirolimus (**Fig. S2G**). Together, these data indicate that mTORC1 activation also occurs in cancer cells upon p16 knockdown, which correlates with increased sensitivity to the mTORC1 inhibitor temsirolimus.

Finally, we aimed to determine whether increased mTORC1 signaling is dependent on RB. While knockdown of RB suppressed senescence (**Fig. S2H-K**), it did not increase p-S6K or p-4EBP1 in either shRRM2-nor BRAF^V600E^-expressing cells (**Fig. S2L-M**). Similarly, knockdown of RB in cancer cell lines did not increase mTORC1 signaling (**Fig. S2N**). Consistently, terms associated with mTOR signaling were not observed using a publicly-available dataset of RB knockdown in senescence (**Table S6**) (Chicas et al., 2010). This suggests that the upregulation of mTORC1 activity is due to an RB-independent pathway downstream of p16 loss. Importantly, these results also indicate that the increase in mTORC1 activity is not a cell cycle-dependent phenomenon. Together, these data demonstrate that activation of mTORC1 signaling downstream of p16 suppression is critical for nucleotide synthesis in an RB-independent manner.

### mTORC1 activation by p16 knockdown increases translation of ribose-5-phosphate isomerase A and promotes nucleotide synthesis through the pentose phosphate pathway

mTORC1 was activated downstream of p16 knockdown to increase nucleotide synthesis (**Fig. 2**). Previous reports have shown that mTORC1 upregulates purine and pyrimidine metabolism through ATF4-*MTHFD2* and CAD, respectively (Ben-Sahra et al., 2013; Ben-Sahra et al., 2016). We did not observe an increase in *MTHFD2* transcription or CAD phosphorylation between shRRM2 and shRRM2/shp16 (**Fig. S3A-B**). These data suggest that an alternative mechanism is regulating nucleotide synthesis downstream of mTORC1 in our model. mTORC1 activity increases translation (Ma and Blenis, 2009); therefore, we aimed to determine whether the observed increase in mTORC1-mediated nucleotide synthesis upon p16 suppression increases translation of transcripts involved in nucleotide synthesis. Towards this goal, we performed polysome fractionation (**Fig. S3C**) followed by RT-qPCR analysis of transcripts involved in purine and pyrimidine synthesis and related anaplerotic pathways (**Table S7**). The positive control *EEF2* (Thoreen et al., 2012) was increased in the heavy polysome fraction compared to the light fraction in shRRM2/shp16 cells (**Fig. S3D**), furthering supporting the notion of increased mTORC1 activity in these cells. Our results reveal a number of transcripts whose abundance is upregulated in the heavy polysome fraction and downregulated in the light polysome fraction (**Fig. 3A and Table S7**), suggesting that these are translationally upregulated upon suppression of p16. We decided to focus only on those transcripts that were significantly upregulated in the heavy fraction and downregulated in the light fraction in shRRM2/shp16 cells. Additionally, both purines and pyrimidines were increased in shRRM2/shp16 cells (i.e., **Fig. 1K-N**). Thus, we further narrowed the list to those transcripts that play an important role in both purine and pyrimidine nucleotide synthesis. Using these criteria, we narrowed the list down to three “hits”: ribose-5-phosphate isomerase A (*RPIA*), nucleoside diphosphate kinase A (*NME1*), and nucleoside diphosphate kinase 3 (*NME3*) (**Fig. 3A**). NME1/NME3 are known metastasis suppressors (Boissan et al., 2018); therefore, we focused on *RPIA*. RPIA is an enzyme that catalyzes the first step of the non-oxidative branch of the pentose phosphate pathway (PPP), which is critical for forming the ribose sugar backbone of both purine and pyrimidine nucleotides (Lane and Fan, 2015). While MYC has been previously shown to increase *RPIA* transcription (Santana-Codina et al., 2018), we did not observe changes in *RPIA* gene expression (**Fig. 3B**) or MYC protein expression in these cells (**Fig. S3E**). Consistent with the idea that mTORC1 regulates *RPIA* translation, RPIA protein expression was increased after p16 suppression and decreased upon mTORC1 inhibition with temsirolimus in both shRRM2/shp16 and BRAF^V600E^/shp16 cells (**Fig. 3C-D**). Increased RPIA protein expression was also observed in our panel of seven isogenic cell lines upon p16 suppression (**Fig. 3E-F**). To further validate these results using a genetic approach, knockdown of regulatory associated protein of MTOR complex 1 (*RPTOR*) decreased RPIA protein expression (**Fig. S3F**). Inhibition of mTORC1 using temsirolimus shifted the *RPIA* mRNA from the heavy to the light polysome fraction (**Fig. 3G**) while it did not decrease total *RPIA* mRNA expression (**Fig. S3G**), further suggesting a role for mTORC1-mediated translation of *RPIA*. Next we aimed to confirm that mTORC1 is directly affecting *RPIA* translation. Previous publications have demonstrated that a short treatment with Torin 1 is a suitable model to study the direct translational targets of mTORC1 (Thoreen et al., 2012). Treatment of shp16 cells for 3 hours with Torin 1 decreased RPIA protein expression and significantly shifted *RPIA* mRNA from the heavy to the light fraction in multiple cell lines (**Fig. 3H-I**). Additionally, knockdown of p16 increased 35S-methionine/cysteine incorporation into RPIA, which was decreased by Torin 1 treatment (**Fig. 3J**). Together, these data demonstrate that mTORC1 directly regulates translation of *RPIA* downstream of p16 knockdown.

**Figure 3.**
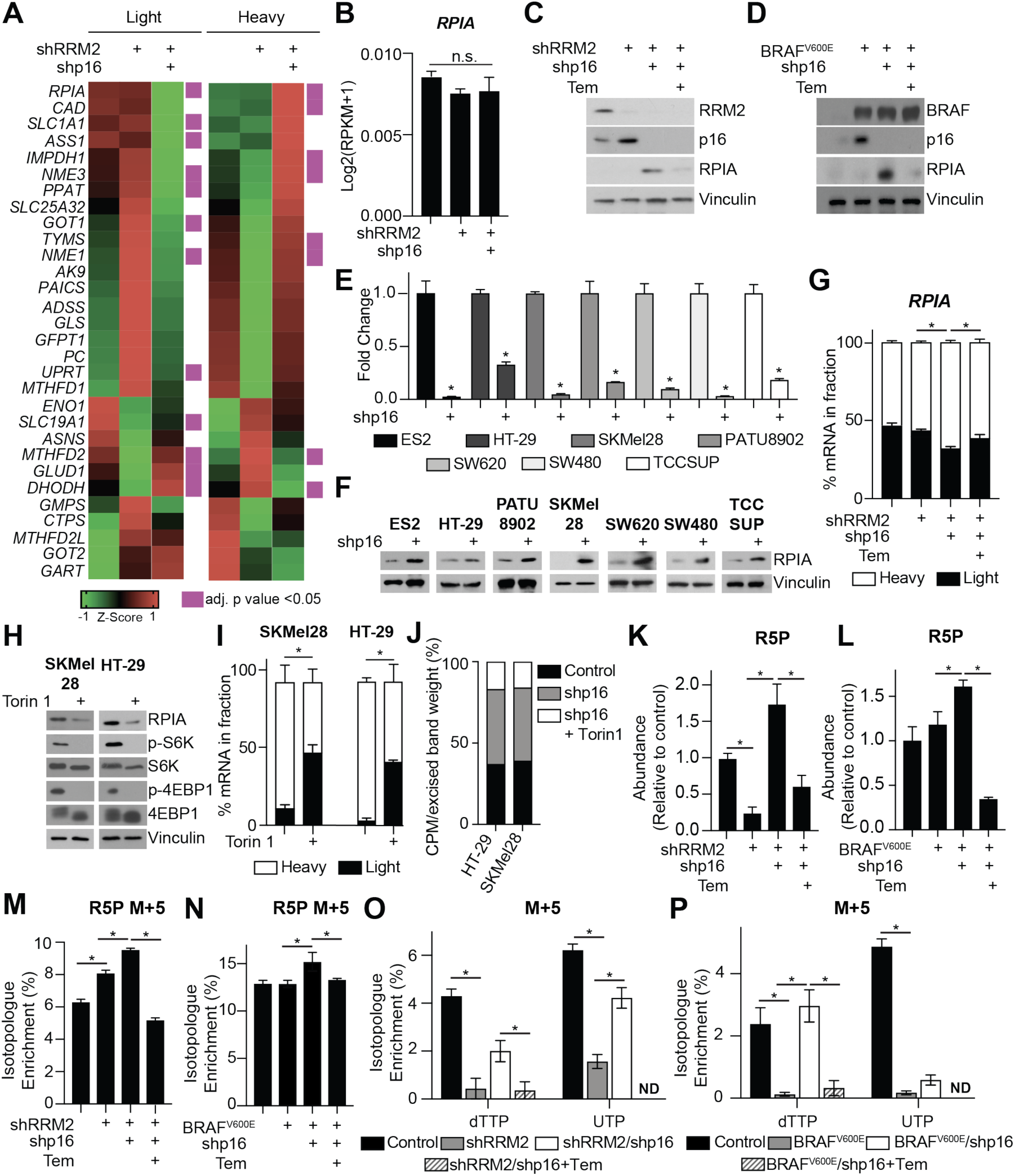
Suppression of p16 increases mTORC1-mediated *RPIA* translation and nucleotide synthesis via the pentose phosphate pathway. **(A-B)** Normal diploid IMR90 cells were infected with lentivirus expressing short hairpin RNA (shRNAs) targeting RRM2 (shRRM2) alone or in combination with an shRNA targeting p16 (shp16). Empty vector was used as a control. **(A)** Heatmap of light and heavy fractions from polysome profiling (see **Table S7** for raw data). This experiment was performed once with 3 technical replicates. **(B)** Total *RPIA* expression from RNA-Seq. Data represent mean ± SEM. ns=not significant. **(C)** Same as (A) but temsirolimus (Tem; 0.5 nM) was added 4 days after starting selection. Immunoblot analysis of the indicated proteins. One of 3 experiments is shown. **(D)** Normal diploid IMR90 cells were infected with lentivirus expressing BRAF^V600E^ alone or in combination with shp16. Empty vector was used as a control. Temsirolimus (Tem; 0.5 nM) was added 4 days after starting selection. Immunoblot analysis of the indicated proteins. One of 3 experiments is shown. **(E)** The indicated cancer cell lines with wildtype p16 expression were infected with a short hairpin targeting p16 and *CDKN2A* expression was determined by RT-qPCR. *p<0.006 **(F)** Same as (E) but cells were serum starved for 16h after which they were incubated with 10% FBS for 30 min. Immunoblot analysis of RPIA. One of at least 2 experiments is shown. **(G)** Percentage of *RPIA* mRNA abundance in polysome fractions in the indicated conditions. *p<0.05 **(H-J)** SKMel28 and HT-29 cells were infected with lentivirus expressing shp16. Cells were serum starved for 16h after which they were incubated with Torin 1 (250nM) in 0.5% FBS for 3 h. **(H)** Immunoblot analysis of the indicated proteins **(I)** Percentage of *RPIA* mRNA abundance in polysome fractions. **(J)** Cells were incubated for 30min with [35S]-methionine/cysteine. RPIA was immunoprecipitated, resolved by SDS-PAGE, and counts per million (CPM) were assessed by scintillation counting. Data represent the percentage of CPM normalized by the excised band weight in the indicated conditions. One of 2 experiments is shown. **(K-L)** Ribose-5-phosphate (R5P) abundance was determined by LC-HRMS. n>3/group, one of at least 3 experiments is shown. Data represent mean ± SEM. *p<0.01 **(M-N)** Cells were incubated with U-^13^C glucose for 8 hours. R5P M+5 was detected in the indicated groups by LC-HRMS. n>3/group, one of at least 3 experiments is shown. Data represent mean ± SEM. *p<0.05 **(O-P)** cells were incubated with U-^13^C glucose for 8 hours. dTTP M+5 and UTP M+5 was detected in the indicated groups by LC-HRMS. n>3/group, one of at least 3 experiments is shown. Data represent mean ± SEM. *p<0.03; ND=not detected

RPIA activity is critical for ribose-5-phosphate (R5P) synthesis. Consistent with increased RPIA protein expression, total R5P was increased in both shRRM2/shp16 and BRAF^V600E^/shp16 cells and decreased by inhibition of mTORC1 (**Fig. 3K-L**). Interestingly, R5P levels were decreased in shRRM2 alone cells, suggesting that upstream metabolic pathways are inhibited at this late time point in this model of senescence. Finally, stable isotope labeling using U-^13^C glucose in both shRRM2/shp16 and BRAF^V600E^/shp16 models demonstrated an increase in the M+5 fraction of R5P and multiple nucleotides upon p16 knockdown, which was abrogated by temsirolimus treatment (**Fig. 3M-P** and **Table S8**). Taken together, these data demonstrate that knockdown of p16 increases mTORC1-mediated translation of RPIA to fuel R5P and nucleotide synthesis.

### Ribose-5-phosphate isomerase A is a metabolic vulnerability of p16-low cells *in vitro* and *in vivo*

RPIA translation and protein expression is increased upon p16 knockdown, which correlated with increased R5P and nucleotide synthesis (**Fig. 3**). To determine whether RPIA is critical for BRAF^V600E^/shp16 or shRRM2/shp16 cell proliferation, we depleted RPIA using two independent shRNAs. Our data indicate that RPIA is necessary for BRAF^V600E^/shp16 and shRRM2/shp16 cell proliferation as shown by increased senescence (**Fig. 4A-E and Fig. S4A-F**). Knockdown of RPIA alone had no effect on parental cells (**Fig. S4A-F**). To determine whether low p16 expression creates a vulnerability to RPIA inhibition in cancer cells, we knocked down p16 and RPIA alone or in combination (**Fig. S5A**). Knockdown of RPIA in combination with p16 knockdown induced cellular senescence as shown by increased cytoplasm, flat morphology, and SA-β-Gal activity, cell cycle arrest, and decreased *CCNA2* and *LMNB1* (**Fig. 4F-I and S5B-F**). TCCSUP cells had the most robust increase in SA-β-Gal activity (**Fig. S5E**), which may be due to the fact that RPIA knockdown was especially robust in these cells (**Fig. S5A**). Knockdown of RPIA alone did not affect cancer cell senescence or proliferation (**Fig. Fig. 4F-I and S5B-F**). We did not observe a marked increase in cell death (**Fig. 5SG**), suggesting that the observed loss of proliferation is likely due to the senescence-associated cell cycle arrest. Similar results were observed *in vivo*, where knockdown of RPIA inhibited tumor growth in HT-29 cells with shp16 but not controls (**Fig. 4J-L**). Consistent with our *in vitro* data, *LMNB1* was decreased only in shp16/shRPIA tumors (**Fig. 4M**). While there was a decrease in *CCNA2* upon RPIA knockdown alone, the difference was significantly larger in shp16/shRPIA compared to shp16 alone tumors (**Fig. 4N**). Together, these data indicate that RPIA-mediated increased nucleotide synthesis is necessary for cancer cell proliferation and that suppression of RPIA may be a target for cancers with low p16 expression.

**Figure 4.**
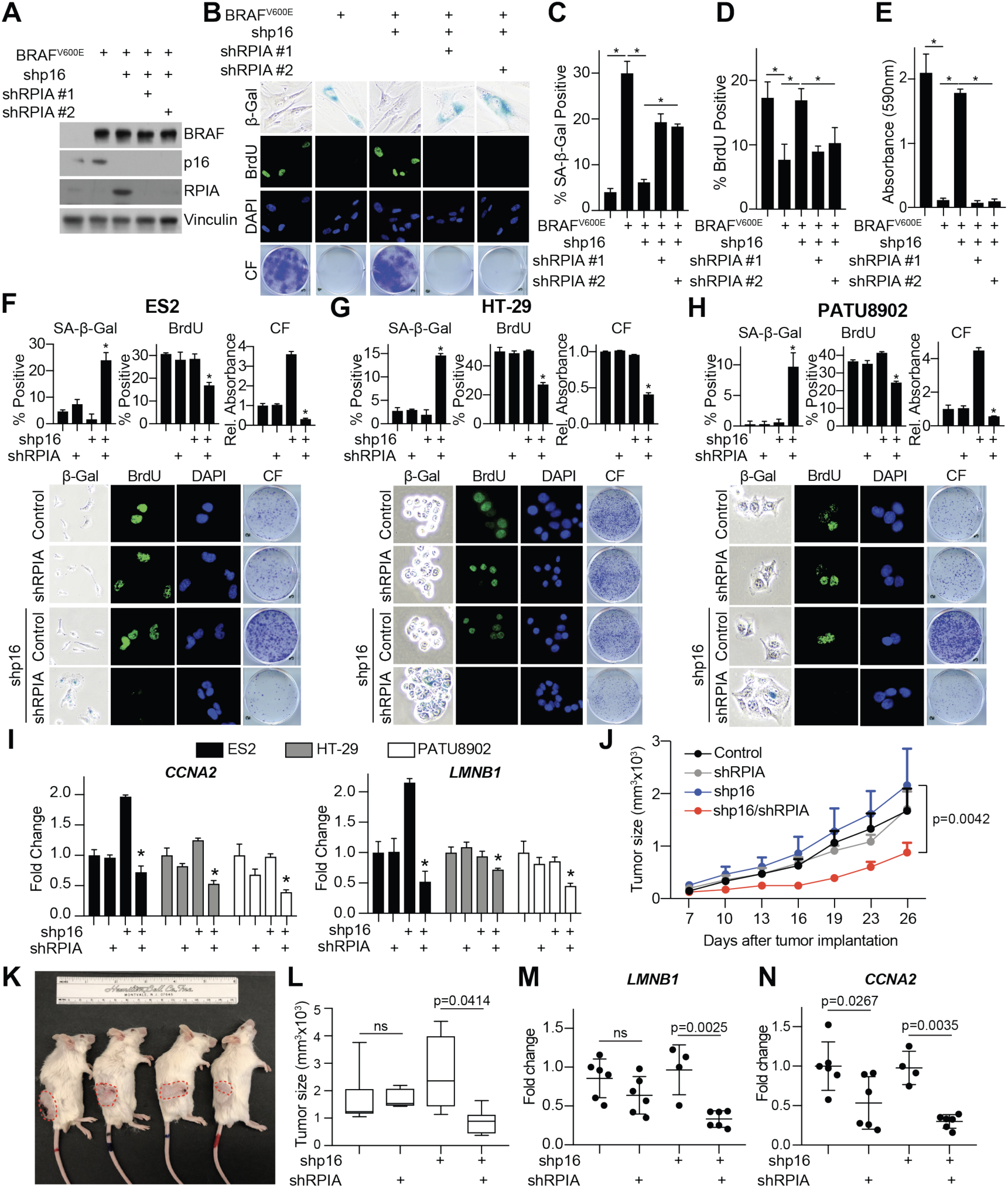
Inhibition of RPIA is a metabolic vulnerability for cells with low p16. **(A-E)** Normal diploid IMR90 cells were infected with retrovirus expressing BRAF^V600E^ alone or in combination with 2 independent shRNAs targeting RPIA (shRPIA). Empty vector was used as a control. **(A)** Immunoblot analysis of the indicated proteins. One of 3 experiments is shown. **(B)** SA-β-Gal activity, BrdU incorporation, and colony formation. One of 3 experiments is shown. **(C)** Quantification of SA-β-Gal activity in (B). n=3/group, one of 3 experiments is shown. Data represent mean ± SEM. *p<0.001 **(D)** Quantification of BrdU incorporation in (B). n=3/group, one of 3 experiments is shown. Data represent mean ± SEM. *p<0.02 **(E)** Quantification of colony formation in (B). n=3/group, one of 3 experiments is shown. Data represent mean ± SEM. *p<0.001 **(F-H)** The indicated cancer cell lines with wildtype p16 expression were infected with a short hairpin targeting RPIA alone or in combination with a shRNA targeting p16. SA-β-Gal activity, BrdU incorporation, and colony formation (CF) for each of the indicated cell lines. n=3/group, one of at least 2 experiments is shown. Data represent mean ± SEM. *p<0.05 vs. shp16 alone. **(I)** *CCNA2* and *LMNB1* fold change in the indicated cells. One of at least 2 experiments is shown. Data represent mean ± SD. *p<0.05 vs. shp16 alone. **(J)** HT-29 colon cancer cells expressing control or p16 shRNA alone or in combination with RPIA shRNA were injected into the flank of SCID mice (4-6 mice/group). Shown is the tumor growth curve over 26 days. **(K)** Representative images of mice from each group. Tumors are outlined in red. **(L)** Tumor volume at Day 26 post-implantation. *p=0.0414; ns= not significant **(M)** *LMNB1* expression was determined by RT-qPCR in the indicated tumors. ns = not significant **(N)** *CCNA2* expression was determined by RT-qPCR in the indicated tumors.

## Discussion

The absence of p16 predisposes cells to tumorigenesis (LaPak and Burd, 2014), and its expression is low or null in many human cancers (Cerami et al., 2012; Gao et al., 2013). There is currently no approved targeted therapy for p16 low tumors (Otto and Sicinski, 2017). Therefore, delineating the molecular mechanisms downstream of p16 suppression is critical to identifying new therapeutics for these patients. While the role of p16 loss in deregulating the cell cycle has been known for decades (Sherr, 2001), its role in metabolism is unclear. In this study, we found that mTORC1 signaling activation upon p16 suppression increases nucleotide synthesis. Mechanistically, we found mTORC1 activity led to increased translation of *RPIA* and glucose flux through the PPP to increase nucleotide levels. Suppression of p16 in cancer cells also leads to increased mTORC1 activity and increased RPIA protein expression, and these cells are more sensitive to both mTORC1 inhibitors or RPIA suppression than p16 wildtype cells. Together, our results suggest that nucleotide metabolism via RPIA is a metabolic vulnerability of p16-null cancers.

Metabolic reprogramming is a hallmark of cancer (Hanahan and Weinberg, 2011). Cancer cells reprogram metabolism to increase biomass needed for growth and proliferation (Pavlova and Thompson, 2016). Modulation of nucleotide and deoxyribonucleotide levels is critical for multiple cancer cell phenotypes, including to repair damaged DNA and ensure rapid proliferation (Kohnken et al., 2015). We previously found that increased deoxyribonucleotides, either through upregulation of RRM2 expression or loss of ATM, bypasses senescence (Aird et al., 2015; Aird et al., 2013). Additionally, a recent paper found that metabolic reprogramming, including increased nucleotide levels, precedes tumor formation in a UVB-induced skin cancer model (Hosseini et al., 2018). Here, we show for the first time that loss of p16 increases nucleotide synthesis (**Fig. 1**) through a mechanism mediated by mTORC1 (**Fig. 3**). Excitingly, activation of this pathway increased both nucleotides and deoxyribonucleotides. We observed an increase in the other ribonucleotide reductase R2 subunit RRM2B (**Fig. S1L-M**). RRM2B has been shown to play a role in mitochondrial dNTP synthesis and in response to DNA damage (Bourdon et al., 2007; Pontarin et al., 2012). Interestingly, RRM2B was increased in both shRRM2 alone and shRRM2/shp16 cells. This suggests that while RRM2B is likely important for reducing NDPs/NTPs to dNDPs/dNTPs in senescence bypass, its upregulation alone is not sufficient to produce dNDPs/dNTPs. Indeed, these data further support that notion that it is only when upstream nucleotides are also increased, such as when p16 is knocked down (**Fig. 1**), that the expression of RRM2B is critical for senescence bypass.

The canonical function of p16 is upstream of RB to affect E2F and the cell cycle (Sherr, 2001). We found that mTORC1 upregulation downstream of p16 knockdown is independent of RB in multiple cell types (**Fig. S2**). There are an increasing number of studies reporting RB-independent functions of p16 (Al-Khalaf et al., 2013; Jenkins et al., 2011; Lee et al., 2013; Tyagi et al., 2017), suggesting that the non-canonical pathway of p16 loss needs to be explored to identify both mechanistic underpinnings of RB-independent functions and novel therapies for cancer patients with p16-null tumors. As knockdown of p16 increased mTORC at the lysosomal membrane (**Fig. S2A**), it is possible that this pathway affects amino acid transporters and/or uptake. Future studies will determine both transcriptional activation of amino acid transporters and amino acid abundance in cells with p16 loss compared to RB loss. Additionally, a previous report in a mouse model of melanomagenesis also found increased mTORC1 signaling upon *Cdkn2a* knockout due to miR-99/100 expression (Damsky et al., 2015). Therefore, it is also possible that p16 and RB differentially regulate miRNA expression. Finally, it is possible that the effect of p16 knockdown is through cyclin D1/CDK4, which has previously been shown to phosphorylate TSC2, thereby activating mTORC1 signaling (Goel et al., 2016). Further research is needed to understand the connection between loss of p16 and the activation of mTORC1.

mTORC1 is a master regulator of metabolism by coordinating metabolite availability though translational control of metabolic enzymes (Iurlaro et al., 2014; Zoncu et al., 2011). Recent studies have linked mTORC1 to both purine and pyrimidine synthesis via *MTHFD2* or CAD, respectively (Ben-Sahra et al., 2013; Ben-Sahra et al., 2016). However, we did not observe an increase in either pathway in shRRM2/shp16 cells. Instead, our results indicate that suppression of p16 increases mTORC1-mediated translation of RPIA. Previous studies have shown that *RPIA* is transcriptionally regulated by mTORC1 signaling (Duvel et al., 2010) or MYC (Santana-Codina et al., 2018). We did not observe a transcriptional increase in *RPIA* or MYC upregulation in our model (**Fig. 3B and S3E**), suggesting that increased *RPIA* upregulation is context-dependent. Consistent with our results, a recent paper showed that total *RPIA* mRNA expression is not significantly decreased even after 24 hours of Torin 1 treatment (Park et al., 2017). Instead, our data demonstrate that RPIA is directly translationally regulated by mTORC1 as inhibition of mTORC1 with a short Torin 1 treatment decreased *RPIA* transcripts in the heavy polysome fraction as well as RPIA protein expression (**Fig. 3**). Consistent with the idea that our results are MYC-independent, a previous publication demonstrated that *MYC* mRNA is resistant to Torin 1 inhibition (Thoreen et al., 2012). Taken together, these data demonstrate that the observed increase in RPIA protein upon loss of p16 is mediated though mTORC1 specific translation. mTORC1 directly mediates translation of mRNAs through terminal oligopyrimidine motif (TOP) sequences, TOP-like sequences, or specific types of 5’UTRs (Gandin et al., 2016; Thoreen et al., 2012). *RPIA* has a putative TOP-like sequence at one of its predicted transcription start sites, suggesting the mTORC1 may regulate *RPIA* translation via this motif. Future studies are required to determine whether mTORC1 is directly regulating *RPIA* translation through a TOP-like sequence.

Cell cycle inhibitors are currently being tested in the clinic for tumors with deletions/mutations in *CDKN2A* (clinicaltrials.gov); however, no FDA-approved therapy currently exists for this subset of patients. Moreover, our data and others demonstrate that p16 may have functions outside of the cell cycle and RB (Al-Khalaf et al., 2013; Jenkins et al., 2011; Lee et al., 2013; Tyagi et al., 2017), suggesting that these inhibitors may not be efficacious in these patients. Excitingly, our results with p16 suppression opens up a metabolic vulnerability through activation of mTORC1-mediated nucleotide metabolism. Indeed, we found that isogenic p16-null cells are more sensitive to temsirolimus or suppression of RPIA both *in vitro* and *in vivo* (**Fig. 2 and 4**). RPIA inhibition has been shown to limit the growth of Kras^G12D^ cell lines and xenografted tumors (Santana-Codina et al., 2018; Ying et al., 2012). Our results demonstrate that RPIA expression could also be exploited as a metabolic target in p16-null cancers.

In conclusion, our study provides a new molecular effect of p16 loss whereby mTORC1 signaling is activated to increase nucleotide metabolism. This is different, yet likely linked, to its canonical role in cell cycle regulation. These mechanistic insights have broad implications for understanding pro-tumorigenic metabolism. Moreover, this study provides a new metabolic vulnerability for p16 low cancer cells, which may be exploited for therapy.

## Acknowledgments

We would like to acknowledge Dr. Alice Soragni (UCLA), Dr. Kristin Eckert and Dr. Nadine Hempel (Penn State College of Medicine), Dr. Gina DeNicola (Moffitt Cancer Center), and Dr. Rugang Zhang (The Wistar Institute) for providing cell lines. We would like to thank Drs. Juan Andres Melendez and Robert P. Feehan for providing helpful discussion. This work was supported by grants from the National Institutes of Health (P50CA174523 and U54CA224070 to M.H.; DK13499 and DK15648 to S.R.K.; K22ES026235 to N.W.S.; and R00CA194309 to K.M.A.), the Dr. Miriam and Sheldon G. Adelson Medical Research Foundation (to M.H.), and the W. W. Smith Charitable Trust (to K.M.A.). The RPPA Core Facility is funded by NCI CA16672.

## Author Contributions

Conceptualization, R.B. and K.M.A.; Methodology, R.K., Y.I., N.W.S.; Investigation, R.B., C.W.C, E.S.D., K.E.L., R.K., N.M., M.N., M.D., H.J., M.Z., L.K., H.L., Y.I., N.W.S., K.M.A.; Resources, G.Z., R.G., G.R., J.I.W., M.H., D.J.D.; Writing, R.B., N.W.S, and K.M.A.; Visualization, R.B., C.W.C, R.K., N.S.W., K.M.A.; Supervision, M.H., Y.L., G.B.M., G.R., J.I.W., S.R. K., D.J.D., N.W.S., K.M.A.; Funding Acquisition, G.B.M., M.H., N.W.S., and K.M.A.

## STAR Methods

### Contact for Reagent and Resource Sharing

Further information and requests for resources and reagents should be directed to and will be fulfilled by the Lead Contact, Katherine M. Aird.

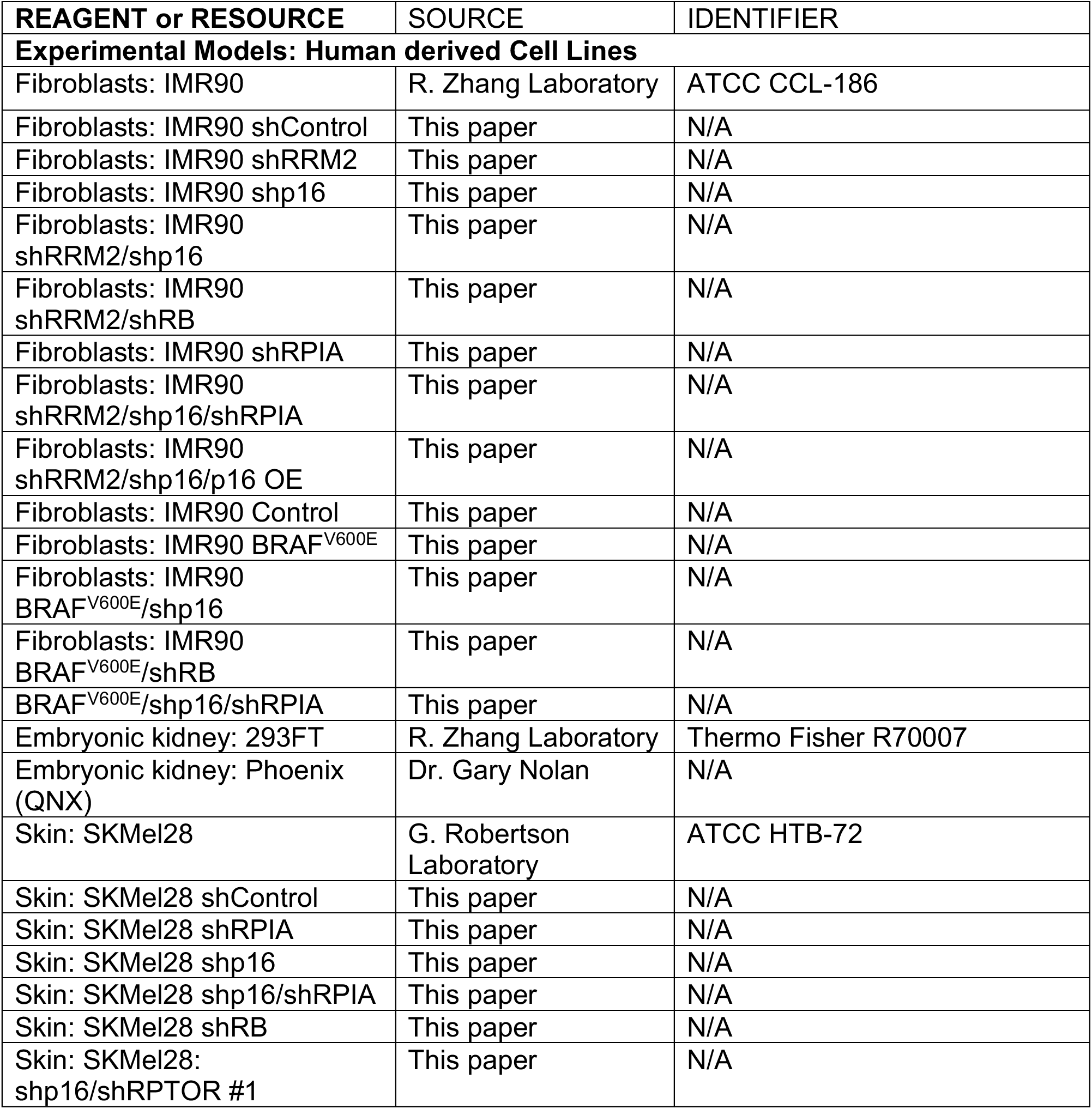
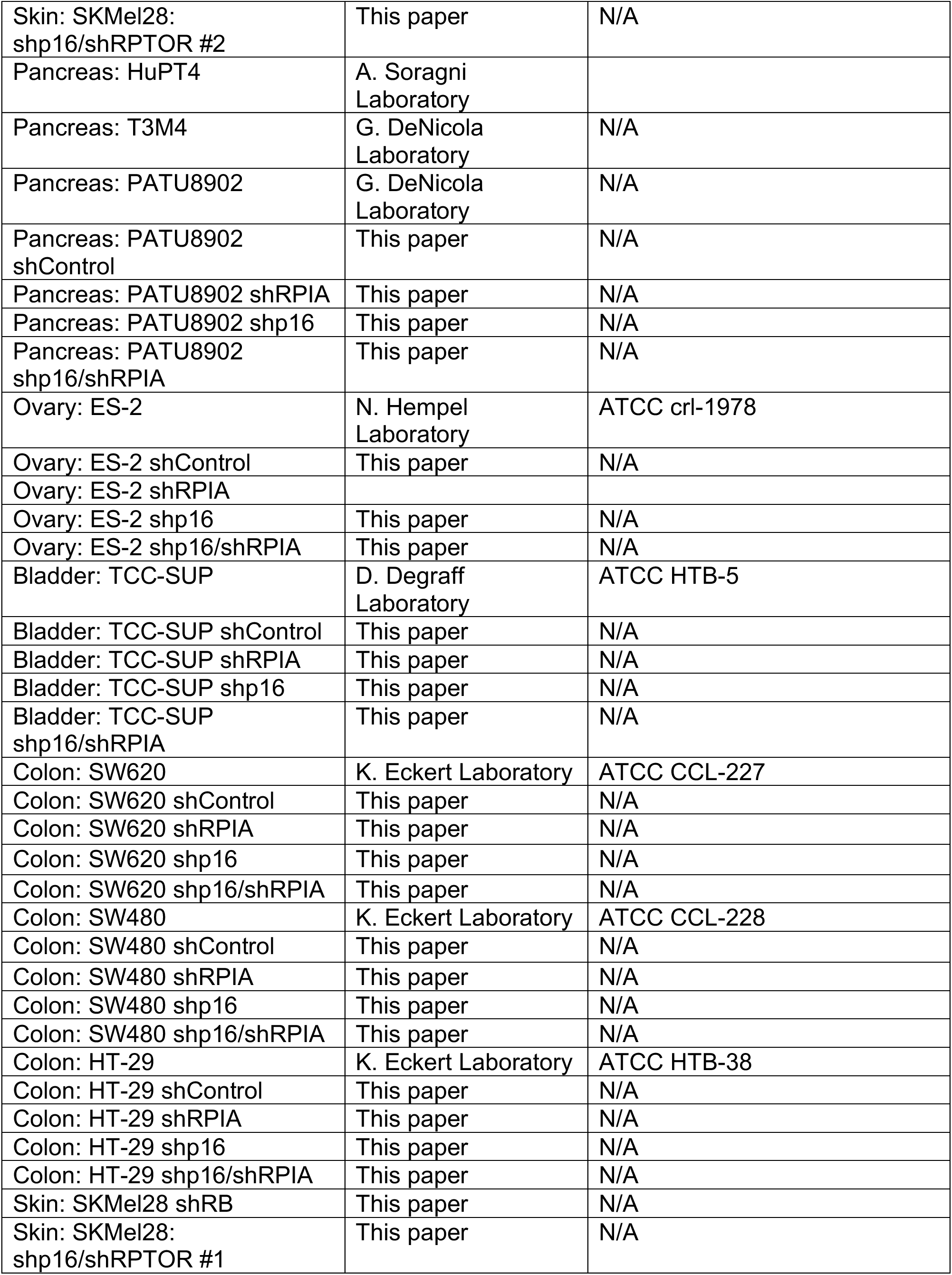
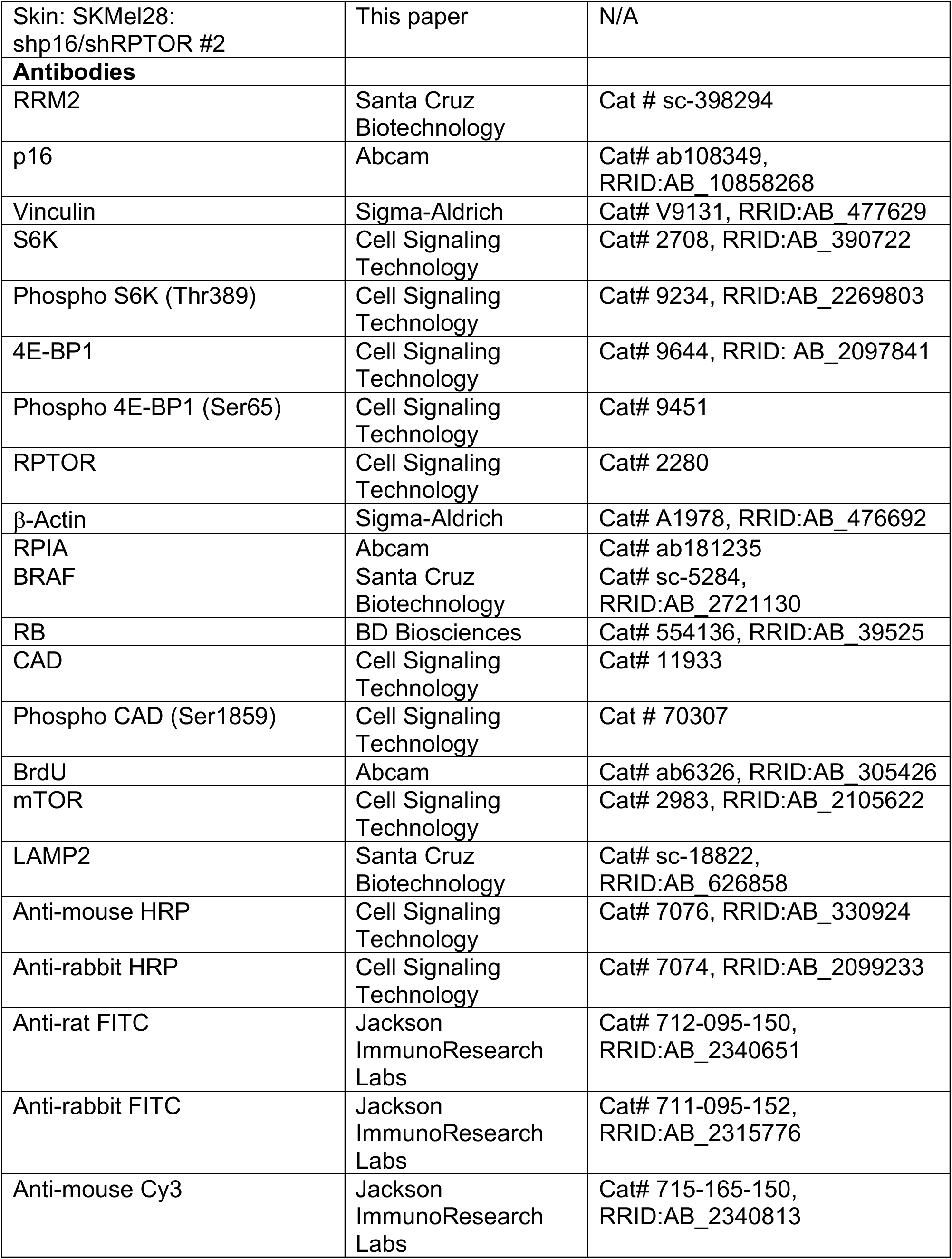
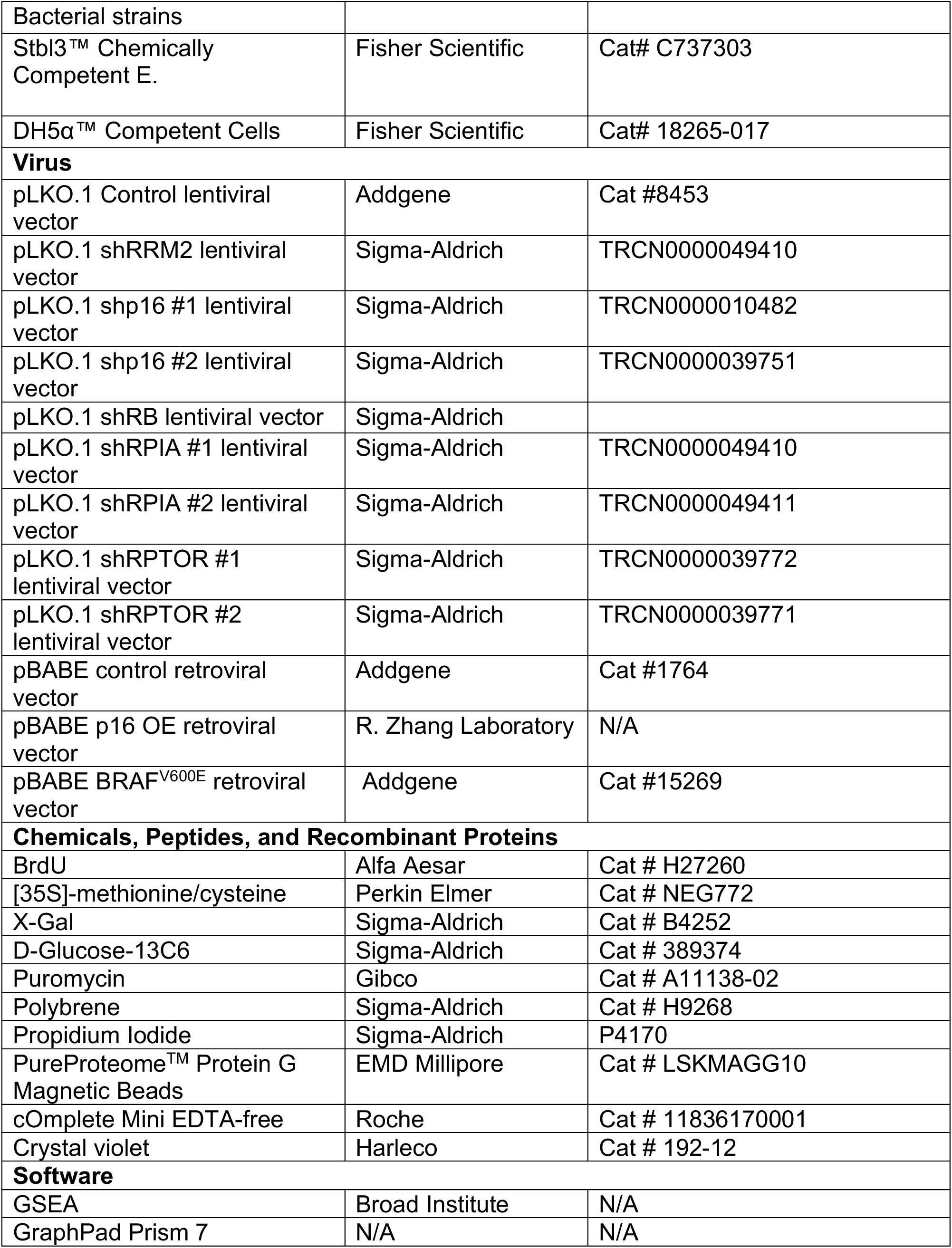
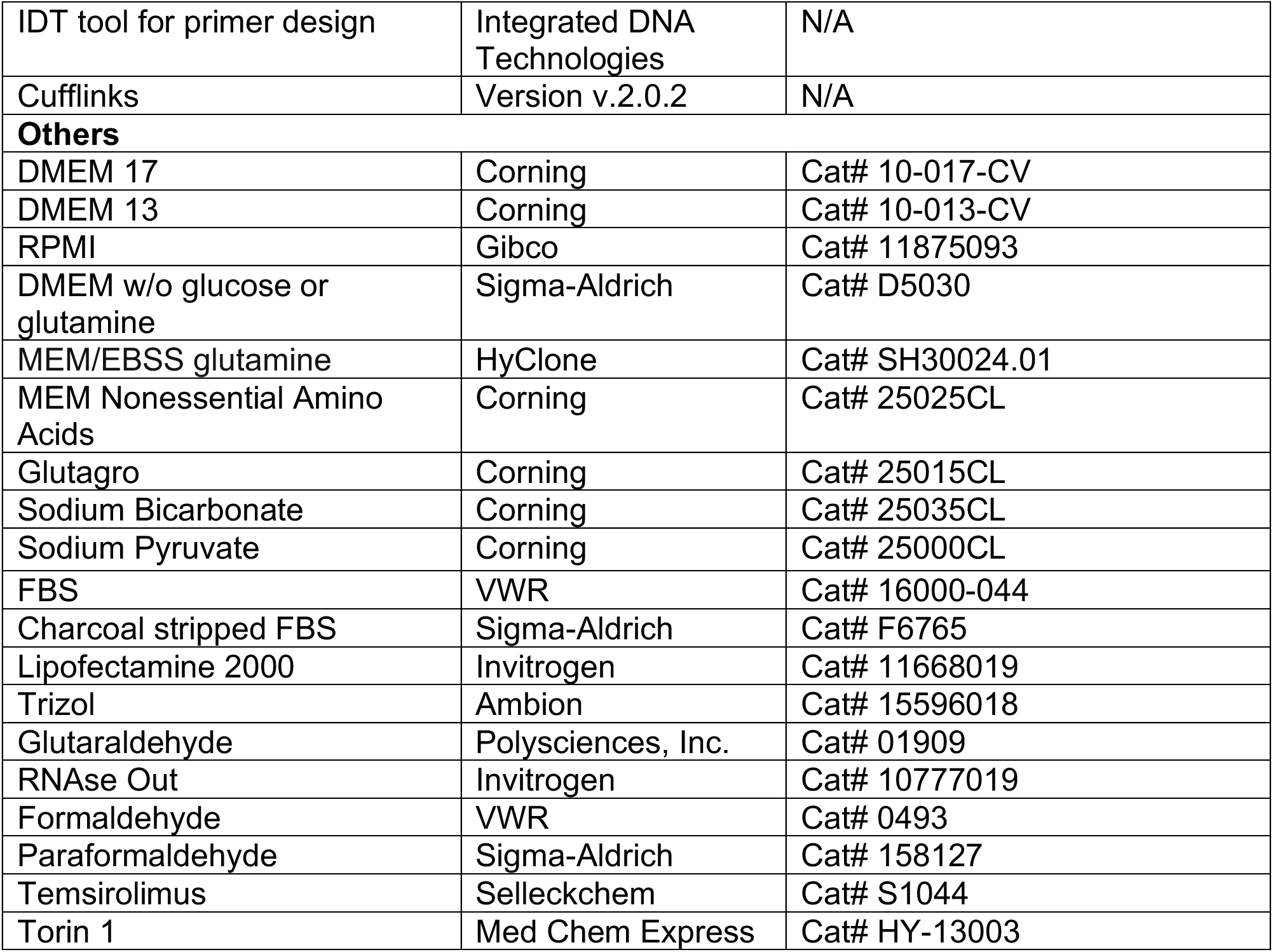

| REAGENT or RESOURCE | SOURCE | IDENTIFIER |
| --- | --- | --- |
| **Experimental Models: Human derived Cell Lines** |  |  |
| Fibroblasts: IMR90 | R. Zhang Laboratory | ATCC CCL-186 |
| Fibroblasts: IMR90 shControl | This paper | N/A |
| Fibroblasts: IMR90 shRRM2 | This paper | N/A |
| Fibroblasts: IMR90 shp16 | This paper | N/A |
| Fibroblasts: IMR90 shRRM2/shp16 | This paper | N/A |
| Fibroblasts: IMR90 shRRM2/shRB | This paper | N/A |
| Fibroblasts: IMR90 shRPIA | This paper | N/A |
| Fibroblasts: IMR90 shRRM2/shp16/shRPIA | This paper | N/A |
| Fibroblasts: IMR90 shRRM2/shp16/p16 OE | This paper | N/A |
| Fibroblasts: IMR90 Control | This paper | N/A |
| Fibroblasts: IMR90 BRAF<sup>V600E</sup> | This paper | N/A |
| Fibroblasts: IMR90 BRAF<sup>V600E</sup>/shp16 | This paper | N/A |
| Fibroblasts: IMR90 BRAF<sup>V600E</sup>/shRB | This paper | N/A |
| BRAF<sup>V600E</sup>/shp16/shRPIA | This paper | N/A |
| Embryonic kidney: 293FT | R. Zhang Laboratory | Thermo Fisher R70007 |
| Embryonic kidney: Phoenix (QNX) | Dr. Gary Nolan | N/A |
| Skin: SKMel28 | G. Robertson Laboratory | ATCC HTB-72 |
| Skin: SKMel28 shControl | This paper | N/A |
| Skin: SKMel28 shRPIA | This paper | N/A |
| Skin: SKMel28 shp16 | This paper | N/A |
| Skin: SKMel28 shp16/shRPIA | This paper | N/A |
| Skin: SKMel28 shRB | This paper | N/A |
| Skin: SKMel28: shp16/shRPTOR #1 | This paper | N/A |
| Skin: SKMel28:<br>shp16/shRPTOR #2 | This paper | N/A |
| Pancreas: HuPT4 | A. Soragni<br>Laboratory |  |
| Pancreas: T3M4 | G. DeNicola<br>Laboratory | N/A |
| Pancreas: PATU8902 | G. DeNicola<br>Laboratory | N/A |
| Pancreas: PATU8902<br>shControl | This paper | N/A |
| Pancreas: PATU8902 shRPIA | This paper | N/A |
| Pancreas: PATU8902 shp16 | This paper | N/A |
| Pancreas: PATU8902<br>shp16/shRPIA | This paper | N/A |
| Ovary: ES-2 | N. Hempel<br>Laboratory | ATCC crl-1978 |
| Ovary: ES-2 shControl | This paper | N/A |
| Ovary: ES-2 shRPIA |  |  |
| Ovary: ES-2 shp16 | This paper | N/A |
| Ovary: ES-2 shp16/shRPIA | This paper | N/A |
| Bladder: TCC-SUP | D. Degraff<br>Laboratory | ATCC HTB-5 |
| Bladder: TCC-SUP shControl | This paper | N/A |
| Bladder: TCC-SUP shRPIA | This paper | N/A |
| Bladder: TCC-SUP shp16 | This paper | N/A |
| Bladder: TCC-SUP<br>shp16/shRPIA | This paper | N/A |
| Colon: SW620 | K. Eckert Laboratory | ATCC CCL-227 |
| Colon: SW620 shControl | This paper | N/A |
| Colon: SW620 shRPIA | This paper | N/A |
| Colon: SW620 shp16 | This paper | N/A |
| Colon: SW620 shp16/shRPIA | This paper | N/A |
| Colon: SW480 | K. Eckert Laboratory | ATCC CCL-228 |
| Colon: SW480 shControl | This paper | N/A |
| Colon: SW480 shRPIA | This paper | N/A |
| Colon: SW480 shp16 | This paper | N/A |
| Colon: SW480 shp16/shRPIA | This paper | N/A |
| Colon: HT-29 | K. Eckert Laboratory | ATCC HTB-38 |
| Colon: HT-29 shControl | This paper | N/A |
| Colon: HT-29 shRPIA | This paper | N/A |
| Colon: HT-29 shp16 | This paper | N/A |
| Colon: HT-29 shp16/shRPIA | This paper | N/A |
| Skin: SKMel28 shRB | This paper | N/A |
| Skin: SKMel28:<br>shp16/shRPTOR #1 | This paper | N/A |
| Skin: SKMel28:<br>shp16/shRPTOR #2 | This paper | N/A |
| **Antibodies** |  |  |
| RRM2 | Santa Cruz<br>Biotechnology | Cat # sc-398294 |
| p16 | Abcam | Cat# ab108349,<br>RRID:AB_10858268 |
| Vinculin | Sigma-Aldrich | Cat# V9131, RRID:AB_477629 |
| S6K | Cell Signaling<br>Technology | Cat# 2708, RRID:AB_390722 |
| Phospho S6K (Thr389) | Cell Signaling<br>Technology | Cat# 9234, RRID:AB_2269803 |
| 4E-BP1 | Cell Signaling<br>Technology | Cat# 9644, RRID: AB_2097841 |
| Phospho 4E-BP1 (Ser65) | Cell Signaling<br>Technology | Cat# 9451 |
| RPTOR | Cell Signaling<br>Technology | Cat# 2280 |
| β-Actin | Sigma-Aldrich | Cat# A1978, RRID:AB_476692 |
| RPIA | Abcam | Cat# ab181235 |
| BRAF | Santa Cruz<br>Biotechnology | Cat# sc-5284,<br>RRID:AB_2721130 |
| RB | BD Biosciences | Cat# 554136, RRID:AB_39525 |
| CAD | Cell Signaling<br>Technology | Cat# 11933 |
| Phospho CAD (Ser1859) | Cell Signaling<br>Technology | Cat # 70307 |
| BrdU | Abcam | Cat# ab6326, RRID:AB_305426 |
| mTOR | Cell Signaling<br>Technology | Cat# 2983, RRID:AB_2105622 |
| LAMP2 | Santa Cruz<br>Biotechnology | Cat# sc-18822,<br>RRID:AB_626858 |
| Anti-mouse HRP | Cell Signaling<br>Technology | Cat# 7076, RRID:AB_330924 |
| Anti-rabbit HRP | Cell Signaling<br>Technology | Cat# 7074, RRID:AB_2099233 |
| Anti-rat FITC | Jackson<br>ImmunoResearch<br>Labs | Cat# 712-095-150,<br>RRID:AB_2340651 |
| Anti-rabbit FITC | Jackson<br>ImmunoResearch<br>Labs | Cat# 711-095-152,<br>RRID:AB_2315776 |
| Anti-mouse Cy3 | Jackson<br>ImmunoResearch<br>Labs | Cat# 715-165-150,<br>RRID:AB_2340813 |
| Bacterial strains |  |  |
| Stbl3™ Chemically Competent E. | Fisher Scientific | Cat# C737303 |
| DH5α™ Competent Cells | Fisher Scientific | Cat# 18265-017 |
| **Virus** |  |  |
| pLKO.1 Control lentiviral vector | Addgene | Cat #8453 |
| pLKO.1 shRRM2 lentiviral vector | Sigma-Aldrich | TRCN0000049410 |
| pLKO.1 shp16 #1 lentiviral vector | Sigma-Aldrich | TRCN0000010482 |
| pLKO.1 shp16 #2 lentiviral vector | Sigma-Aldrich | TRCN0000039751 |
| pLKO.1 shRB lentiviral vector | Sigma-Aldrich |  |
| pLKO.1 shRPIA #1 lentiviral vector | Sigma-Aldrich | TRCN0000049410 |
| pLKO.1 shRPIA #2 lentiviral vector | Sigma-Aldrich | TRCN0000049411 |
| pLKO.1 shRPTOR #1 lentiviral vector | Sigma-Aldrich | TRCN0000039772 |
| pLKO.1 shRPTOR #2 lentiviral vector | Sigma-Aldrich | TRCN0000039771 |
| pBABE control retroviral vector | Addgene | Cat #1764 |
| pBABE p16 OE retroviral vector | R. Zhang Laboratory | N/A |
| pBABE BRAF<sup>V600E</sup> retroviral vector | Addgene | Cat #15269 |
| **Chemicals, Peptides, and Recombinant Proteins** |  |  |
| BrdU | Alfa Aesar | Cat # H27260 |
| [35S]-methionine/cysteine | Perkin Elmer | Cat # NEG772 |
| X-Gal | Sigma-Aldrich | Cat # B4252 |
| D-Glucose-13C6 | Sigma-Aldrich | Cat # 389374 |
| Puromycin | Gibco | Cat # A11138-02 |
| Polybrene | Sigma-Aldrich | Cat # H9268 |
| Propidium Iodide | Sigma-Aldrich | P4170 |
| PureProteome™ Protein G Magnetic Beads | EMD Millipore | Cat # LSKMAGG10 |
| cOmplete Mini EDTA-free | Roche | Cat # 11836170001 |
| Crystal violet | Harleco | Cat # 192-12 |
| **Software** |  |  |
| GSEA | Broad Institute | N/A |
| GraphPad Prism 7 | N/A | N/A |
| IDT tool for primer design | Integrated DNA Technologies | N/A |
| Cufflinks | Version v.2.0.2 | N/A |
| **Others** |  |  |
| DMEM 17 | Corning | Cat# 10-017-CV |
| DMEM 13 | Corning | Cat# 10-013-CV |
| RPMI | Gibco | Cat# 11875093 |
| DMEM w/o glucose or glutamine | Sigma-Aldrich | Cat# D5030 |
| MEM/EBSS glutamine | HyClone | Cat# SH30024.01 |
| MEM Nonessential Amino Acids | Corning | Cat# 25025CL |
| Glutagro | Corning | Cat# 25015CL |
| Sodium Bicarbonate | Corning | Cat# 25035CL |
| Sodium Pyruvate | Corning | Cat# 25000CL |
| FBS | VWR | Cat# 16000-044 |
| Charcoal stripped FBS | Sigma-Aldrich | Cat# F6765 |
| Lipofectamine 2000 | Invitrogen | Cat# 11668019 |
| Trizol | Ambion | Cat# 15596018 |
| Glutaraldehyde | Polysciences, Inc. | Cat# 01909 |
| RNAse Out | Invitrogen | Cat# 10777019 |
| Formaldehyde | VWR | Cat# 0493 |
| Paraformaldehyde | Sigma-Aldrich | Cat# 158127 |
| Temsirolimus | Selleckchem | Cat# S1044 |
| Torin 1 | Med Chem Express | Cat# HY-13003 |

### Experimental Model and Subject Details

#### Cell Lines

Normal diploid IMR90 human fibroblasts were cultured according to the ATCC in low oxygen (2% O_2_) in DMEM (4.5 g/L glucose, Corning cat# 10017CV) with 10% FBS supplemented with L-glutamine, non-essential amino acids, sodium pyruvate, and sodium bicarbonate. Experiments were performed on IMR90 between population doubling #25-35. Melanoma (SKMel28), pancreatic (PATU8902), colorectal (HT-29, SW620, and SW480,) tumor cells and lentiviral and retroviral packaging cells (293FT and Phoenix, respectively) were cultured in DMEM (Corning, cat# 10013CV) with 10% FBS. ES2 ovary tumor cell line was cultured in RPMI medium 1640 with 10% FBS. TCCSUP bladder cancer cell line was cultured in MEM/EBSS glutamine supplemented with 10% FBS. All cell lines were cultured in MycoZap and were routinely tested for mycoplasma as described in (Uphoff and Drexler, 2005). All cell lines were authenticated using STR Profiling using Genetica DNA Laboratories.

#### Mice

Two-month old male SCID mice were purchased from Charles River Laboratories. All mice were maintained in a HEPA-filtered ventilated rack system at the Milton S. Hershey Medical Center animal facility. Mice were housed up to 4 mice per cage and in a 12-hour light/dark cycle. All experiments with animals were performed in accordance with institutional guidelines approved by the Institutional Animal Care and Use Committee (IACUC) at the Penn State College of Medicine.

### Method details

#### Lentiviral and retroviral packaging and infection

Retrovirus production and transduction were performed using the BBS/calcium chloride method (Aird et al., 2013). Phoenix cells (a gift from Dr. Gary Nolan, Stanford University) were used to package the infection viruses. Lentiviral constructs were transfected into 293FT cells using Lipofectamine 2000 (Thermo Fisher). Lentivirus was packaged using the ViraPower Kit (Invitrogen, Carlsbad, CA, USA) following the manufacturer’s instructions.

The basic IMR90 experiment timeline is delineated in **Figure S1A**. Briefly, IMR90 cells were infected with pLKO.1 empty vector or pLKO.1-shRRM2, and 24 hours later cells were infected with pLKO.1 empty vector, pLKO.1-shp16 or pLKO.1-shRB. Cells were selected with puromycin (3μg/mL) for 7 days. Alternatively, IMR90 cells were infected with pBABE control or pBABE BRAF^V600E^ vector and 24 hours later cells were infected with a second round of pBABE control or BRAF^V600E^ vector together with pLKO.1 empty vector, pLKO.1-shp16 or pLKO.1-shRB. Cells were selected with puromycin (3μg/mL) for 7 days. Where indicated, cells were treated at day 4 with temsirolimus (0.5nM) or infected with pLKO.1-shRPIA. p16 rescue experiment was performed by simultaneous infection with pLKO.1-shp16 and pBABE-p16 overexpression plasmid. For single infections, cells were infected with the corresponding virus and selected in puromycin (1μg/mL) for 7 days.

Tumor cell lines were infected with pLKO.1 empty vector, pLKO.1-shp16 or pLKO.1-shRB. Cells were selected with puromycin (1μg/mL) for 4 days. Where indicated, cells were treated at day 4 with increasing concentrations of temsirolimus (0.07-50μM) or infected with pLKO.1 shRPIA or pLKO.1-shRPTOR. For double infections, cells were selected in puromycin (3μg/mL) for 4 additional days. For Torin 1 experiments, cells were serum starved for 16h and then treated with 250nM Torin 1 for 3h in 0.5% FBS.

#### RNA-Sequencing and Analysis

Total RNA was extracted from cells at day 7 (**Fig. S1A**) with Trizol (Life Technologies) and DNAse treated with RNeasy Mini Kit (Qiagen, cat#74104) following the manufacturer’s instructions. Three technical replicates were used for each sample RNA integrity number (RIN) was measured using BioAnalyzer (Agilent Technologies) RNA 6000 Nano Kit to confirm RIN above 7. The cDNA libraries were prepared using KAPA Stranded RNA-Seq Kits with RiboErase (Kapa Biosystems). Next generation sequencing was performed in The Penn State College of Medicine Genome Sciences and Bioinformatics Core facility as previously described in (Lynch et al., 2015) using a HiSeq 2500 sequencer (Illumina). Demultiplexed and quality-filtered mRNA-Seq reads were then aligned to human reference genome (GRCh38) using TopHat (v.2.0.9). Differential expression analysis was done using Cuffdiff tool which is available by Cufflinks (v.2.0.2) as described in (Lynch et al., 2015). Data are deposited on GEO (GSE133660).

#### Reverse Phase Protein Array (RPPA) performance and analysis

Following the indicated procedure described above, cells cultured in 10cm dishes were incubated on ice with 300uL of lysis buffer (1% Triton X-100, 50mM HEPES pH=7.5, 150mM NaCl, 1.5mM MgCl2, 1mM EGTA, 100mM NaF, 10mM Na pyrophosphate, 1mM Na3VO4 and 10% glycerol) for 20 min with occasional shaking every 5 min. After incubation, cells were scraped off the plate and centrifuged at 14000 rpm for 10 min at 4°C. Total protein was quantified with Bradford assay and 90ug of protein was diluted 3:1 in SDS sample buffer (40% glycerol, 8% SDS, 0.25M Tris-HCl and 10% B-mercaptoethanol). Lysates were boiled at 95°C for 5 min and stored at −80°C. RPPA data was generated and analyzed by the CCSG-supported RPPA Core Facility at the University of Texas MD Anderson Cancer Center (Akbani et al., 2014). A total of 240 authenticated Antibodies for total protein expression and 64 antibodies for protein phosphorylation were analyzed in this study. The complete antibody list can be found in https://goo.gl/XKsv6s.

#### Gene set enrichment analysis (GSEA)

Expression values included in the Talantov data set (18 nevus and 45 primary melanoma tumors) were downloaded from GSE3189, while the expression values included in the Kabbarah data set (9 nevus and 31 primary melanoma tumors) were downloaded from GSE46517. Gene Cluster Text files (GTC), as well as Categorical Class files (CLS) were generated independently for, RPPA, Talantov and Kabbarah data sets following the Gene Set Enrichment Analysis (GSEA) documentation indications (http://software.broadinstitute.org/gsea/index.jsp). GTC and CLS files were used to run independent GSEA analysis (javaGSEA desktop application). GSEA for Hallmarks, KEGG and Reactome were run independently under the following parameters: 1000 permutations, weighted enrichment analysis, signal to noise metric for ranking genes, and “meandiv” normalization mode. Following GSEA documentation indications, terms with p-value ≤ 0.05 and a q-value ≤ 0.25 were considered significant (Table S1 and S3). Genes were ranked according to the fold-change and p-value obtained on the differential gene expression analysis as described in (Plaisier et al., 2010). Pre-ranked files were built up for the RNA-Seq and for Chicas et al. data set (Chicas et al., 2010) and used to run pre-ranked GSEA (Subramanian et al., 2005) under predefined parameters following GSEA documentation indications.

#### Polysome fractionation

Eight culture plates per condition (~23 million cells per condition) were incubated with harringtonine (2μg/mL) for 2 min at 37°C followed by 5 min of cycloheximide (100μg/mL) treatment at 37°C. Cells were washed twice with PBS after each treatment. Cells were scraped in 600uL of lysis buffer (50mM HEPES, 75mM KCl, 5mM MgCl2, 250mM sucrose, 0.1mg/mL cycloheximide, 2mM DTT, 1% Triton X-100 and 1.3% sodium deoxycholate and 5μl of RNase OUT) on ice. Lysates were rocked for 10 min at 4°C and centrifuged at 3000g for 15 min at 4°C. 400μl of lysates supernatant (cytosolic cell extracts) were layered over cold sucrose gradients (10mM HEPES, 75mM KCl, 5mM MgCl2, 0.5mM EDTA and increasing sucrose concentrations from 20% to 47%). Gradients were centrifuged at 34,000 rpms in a Beckman SW41 rotor for 2h and 40 min at 4°C. After centrifugation, low (0 to 2 ribosomes) and high (<2 ribosomes) polysome fractions were collected in Trizol (1:1) using a density gradient fractionation system (Brandel) equipped with a UA-6 absorbance detector and a R1 fraction collector.

#### RT-qPCR

Total RNA was extracted from cells with Trizol, DNase treated, cleaned, and concentrated using Zymo columns (Zymo Research, Cat# R1013) following the manufacturer’s instructions. Similarly, mRNA from polysome fractions was DNase treated, cleaned, and concentrated using Zymo columns. Optical density values of RNA were measured using NanoDrop One (Thermo Scientific) to confirm an A260 and A280 ratio above 1.9. Relative expression of target genes (listed in **Table S9**) were analyzed using the QuantStudio 3 Real-Time PCR System (Thermo Fisher Scientific) with clear 96 well plates (Greiner Bio-One). Primers were designed using the Integrated DNA Technologies (IDT) tool (http://eu.idtdna.com/scitools/Applications/RealTimePCR/) (**Table S9**). A total of 25ng of RNA was used for One-Step qPCR (Quanta BioSciences) following the manufacturer’s instructions in a final volume of 10μl. Conditions for amplification were: 10 min at 48°C, 5 min at 95°C, 40 cycles of 10 s at 95°C and 7 s at the corresponding annealing temperature (**Table S9**). The assay ended with a melting curve program: 15 s at 95°C, 1 min at 70°C, then ramping to 95°C while continuously monitoring fluorescence. Alternatively, relative expression of *CDKN2A* was determined by adapting the method of Zhang Q, et al. (Zhang et al., 2015). Conditions for amplification were: 10 min at 48°C, 5 min at 95°C, 4 cycles of 10s at 95°C and 10s starting at 66°C and decreasing 2°C per cycle, 40 cycles of 10 s at 95°C and 7 s at 64°C. The assay ended with a melting curve program as above. Each sample was assessed in triplicate. Relative quantification was determined to multiple reference genes (*B2M*, *MRPL9*, *PSMC4*, and *PUM1*) using the delta-delta Ct method. The percentage of target gene mRNA in each polysome fraction was calculated similar to (Panda et al., 2017), but Ct values were first normalized to the reference gene *PUM1*.

#### [35S]-methionine/cysteine incorporation followed by IP

After 16h of FBS starvation, cells were treated with 250nM Torin 1 or DMSO in media supplemented with 0.5% FBS for 3h. Thirty minutes prior to the end of Torin 1 treatment, 110uCi [35S]-methionine/cysteine was added to each plate. Cells were wash twice with PBS and pelleted. The pellet was resuspended in 100uL of denaturing Lysis buffer (1mM SDS and 5mM EDTA pH = 8) and boiled at 95°C for 5 min. Lysates were resuspended in 900uL of non-denaturing buffer (20mM Tris-HCl pH = 8, 137mM NaCl, 1mM EDTA pH = 8 and 1% Triton-X), sonicated, and centrifuged (10 min, 13,000g, 4°C). The supernatant was collected, and the protein concentration was determined using the Bradford assay. 500ug of protein per condition was precleared with 15uL of magnetic beads (rotating 1h, 4°C). 5uL per sample of anti-RPIA or anti-IgG were bound to magnetic beads (rotating 3h, 4°C). Precleared samples were incubated with corresponding magnetic beads with conjugated antibodies (rotating overnight, 4°C). Magnetic beads were washed 3 times for 15 min each at 4°C with cold wash buffer (10mM Tris-HCl pH = 7.4, 1mM EDTA pH = 8, 1mM EGTA pH = 8, 150mM NaCl, 1% Triton-X, 0.2mM Na_3_VO_4_, 1mM PMSF and cOmplete Mini EDTA-free 1x). Immunoprecipitates were eluted in 10uL of 1X sample buffer (2% SDS, 10% glycerol, 0.01% bromophenol blue, 62.5mM Tris, pH 6.8, 0.1M DTT) and boiled 10 min at 65°C and 1000rpm. Immunoprecipitates and 10% of input were separated under a 12% acrylamide gel. Acrylamide gel was stained with Coomassie blue and immunoprecipitated bands corresponding to RPIA (35kDa) were excised from the gel. Excised bands were weighted for normalization purposes before digestion with 1ml of electrode buffer (96mM Tris, 500mM glycine and 0.4% w/s SDS) for 16h at 4°C. Excised band suspensions were used to quantify the counts per minute (CPM) in a Beckman LS6500 scintillation counter. CPMs were normalized to corresponding band weights (mg).

#### Senescence and proliferation assays

SA-β-Gal staining was performed as previously described (Dimri et al., 1995). Cells were fixed in 2% formaldehyde/0.2 glutaraldehyde% in PBS (5 min) and stained (40 mM Na_2_HPO_4_, 150 mM NaCl, 2 mM MgCl_2,_ 5 mM K_3_Fe(CN)_6_, 5 mM K_4_Fe(CN)_6_, 1 mg/ml X-gal) overnight at 37°C in a non-CO_2_ incubator. Images were acquired at room temperature using an inverted microscope (Nikon Eclipse Ts2) with a 20X/0.40 objective (Nikon LWD) equipped with a camera (Nikon DS-Fi3). For analysis of SA-β-Gal staining, at least 100 cells per well were counted (>300 cells per experiment).

For BrdU incorporation, cells on coverslips were incubated with 1uM BrdU for 30 min (IMR90, ES2, and SKMel28) or 15 min (SW620, SW480, HT-29, PATU8902, and TCCSUP). Cells were fixed in 4% paraformaldehyde (10min), permeabilized in 0.2% Triton X-100 (5 min), and postfixed in 1% PF 0.01% Tween-20 (30 min). Cells were DNaseI treated (10 min) previous to blocking with 3% BSA/PBS (5 mins). Cells were incubated in anti-BrdU primary antibody in 3% BSA/PBS (1:500) for 1h followed by 1h incubation in FITC anti-Rat secondary antibody in 3% BSA/PBS (1:1000). Finally, cells were incubated with 0.15 μg/ml DAPI in PBS (1 min), mounted, and sealed. Images were acquired at room temperature using a Nikon Eclipse 90i microscope with a 20x/0.17 objective (Nikon DIC N2 Plan Apo) equipped with a CoolSNAP Photometrics camera. For all BrdU experiments, at least 200 cells per coverslip were counted (>600 cells per experiment).

For colony formation, an equal number of cells were seeded in 6-well plates and cultured for an additional 2 weeks. Colony formation was visualized by fixing cells in 1% paraformaldehyde (5 min) and staining with 0.05% crystal violet (20 min). Wells were destained in 500μl 10% acetic acid (5 min). Absorbance (590nm) was measured using a spectrophotometer (Spectra Max 190).

#### Immunofluorescence

Cells were fixed in 4% paraformaldehyde (10min) and permeabilized in 0.2% Triton X-100 (5 min). Cells were blocked with 3% BSA/PBS (5 mins) and incubated in anti-mTORC (1/200) and anti-LAMP2 (1/100) in 3% BSA/PBS (16h). Cells were then incubated in FITC anti-Rabbit (1/2000) and Cy3 anti-mouse (1/5000) secondary antibodies in 3% BSA/PBS (1 hour). for 1 hour followed by 1h incubation in FITC anti-Rat secondary antibody in 3% BSA/PBS (1:1000). Finally, cells were incubated with 0.15 μg/ml DAPI in PBS (1 min), mounted and sealed. Images were acquired at room temperature using a confocal microscope (Leica SP8) with a 64X oil objective. Co-localization analysis was performed using the Leica software.

#### Western blotting

Cell lysates were collected in 1X sample buffer (2% SDS, 10% glycerol, 0.01% bromophenol blue, 62.5mM Tris, pH 6.8, 0.1M DTT) and boiled (10 min at 95°C). Protein concentration was determined using the Bradford assay. Proteins were resolved using SDS-PAGE gels and transferred to nitrocellulose membranes (Fisher Scientific) (110mA for 2 h at 4°C). Membranes were blocked with 5% nonfat milk or 4% BSA in TBS containing 0.1% Tween-20 (TBS-T) for 1 h at room temperature. Membranes were incubated overnight at 4°C in primary antibodies in 4% BSA/TBS + 0.025% sodium azide. Membranes were washed 4 times in TBS-T for 5 min at room temperature after which they were incubated with HRP-conjugated secondary antibodies (Cell Signaling, Danvers, MA) for 1 h at room temperature. After washing 4 times in TBS-T for 5 min at room temperature, proteins were visualized on film after incubation with SuperSignal West Pico PLUS Chemiluminescent Substrate (ThermoFisher, Waltham, MA).

#### Nucleotide Analysis by LC-HRMS

Standards for ADP, dADP, dATP, dTDP, dTTP, CDP, dCDP, CTP, and dCTP and were from Sigma-Aldrich (St Louis, MO). Stable isotope labeled internal standards AMP-^13^C_10_,^15^N_5_, dAMP-^13^C_10_,^15^N_5_, ATP-^13^C_10_,^15^N_5_, dATP-^13^C_10_,^15^N_5_, dTMP-^13^C_10_,^15^N_2_, dTTP-^13^C_10_,^15^N_2_, dCMP-^13^C_9_,^15^N_3_, CTP-^13^C_9_,^15^N_3_, dCTP-^13^C_9_,^15^N_3_, were also from Sigma-Aldrich. No suitable source of stable isotope labeled ADP, dADP, dTDP, GDP, dGDP, CDP, or dCDP was found, thus the mono-phosphate was used as a surrogate internal standard. Diisopropylethylamine (DIPEA) and 1,1,1,3,3,3-hexafluoro 2-propanol (HFIP) were purchased from Sigma-Aldrich. Optima LC-MS grade water, methanol, and acetonitrile (ACN) were purchased from Thermo Fisher Scientific (Waltham, MA).

LC-HRMS for nucleotides and other polar metabolites was as previously described (Guo et al., 2016; Kuskovsky et al., 2019). Briefly, an Ultimate 3000 UHPLC equipped with a refrigerated autosampler (at 6 °C) and a column heater (at 55 °C) with a HSS C18 column (2.1 × 100 mm i.d., 3.5 μm; Waters, Milford, MA) was used for separations. Solvent A was 5 mM DIPEA and 200 mM HFIP and solvent B was methanol with 5 mM DIPEA 200 mM HFIP. The gradient was as follows: 100 % A for 3 min at 0.18 mL/min, 100 % A at 6 min with 0.2 mL/min, 98 % A at 8 min with 0.2 mL/min, 86 % A at 12 min with 0.2 mL/min, 40 % A at 16 min and 1 % A at 17.9 min-18.5 min with 0.3 mL/min then increased to 0.4 mL/min until 20 min. Flow was ramped down to 0.18 mL/min back to 100 % A over a 5 min re-equilibration. For MS analysis, the UHPLC was coupled to a Q Exactive HF mass spectrometer (Thermo Scientific, San Jose, CA, USA) equipped with a HESI II source operating in negative mode. The operating conditions were as follows: spray voltage 4000 V; vaporizer temperature 200 °C; capillary temperature 350 °C; S-lens 60; in-source CID 1.0 eV, resolution 60,000. The sheath gas (nitrogen) and auxiliary gas (nitrogen) pressures were 45 and 10 (arbitrary units), respectively. Single ion monitoring (SIM) windows were acquired around the [M-H]^−^ of each analyte with a 20 *m/z* isolation window, 4 *m/z* isolation window offset, 1e^6^ ACG target and 80 ms IT, alternating in a Full MS scan from 70-950 *m/z* with 1e6 ACG, and 100 ms IT. Data was analyzed in XCalibur v4.0 and/or Tracefinder v4.1 (Thermo) using a 5 ppm window for integration of the peak area of all analytes.

#### Glucose labeling and analysis

Cells were seeded in 10 cm culture plates, and at the end of the indicated treatment media was replaced by 6mL of DMEM (Cat# D5030) supplemented with 0.5% of charcoal stripped FBS, 5mM of ^13^C_6_-D-glucose and 20mM of HEPES. After 8 hours cells were harvested and snap frozen in liquid nitrogen.

Isotopologue patterns for dNDPs, dNTPs and ribose-5-phosphate were analyzed by LC-HRMS as indicated above. Adjustment for natural isotopic abundance was conducted through open source and publicly available FluxFix (Trefely et al., 2016).

#### Flow Cytometry

For 7AAD staining both cells and media were collected and centrifugated (1000 rpm for 5 min) followed by resuspension in 500uL of 7AAD staining solution (5uL of 7AAD solution + 38mM NaCitrate). Stained cells were run on a 10-color FACSCanto flow cytometer (BD biosciences). Data were analyzed with FlowJo Software.

#### Murine tumor model

HT-29 colorectal carcinoma cells were infected with shRNA targeting p16 and RPIA alone or in combination. After 2 days of puromycin selection (3μg/mL), 3 million cells were resuspended in 200μl of PBS and injected subcutaneously into the left flank of SCID mice. Mice were monitored daily to identify palpable tumors. Mice weight and tumor length (L) and width (W) (L>W) were measured every 3 days after a tumor volume of 200mm^3^. Tumor volume was calculated as ½ (L x W^2^). All animals were sacrificed at day 26 post injection and tumor tissues collected for following experiments.

### Quantification and Statistical Analysis

GraphPad Prism version 7.0 was used to perform statistical analysis. One-way ANOVA or t-test were used as appropriate to determine p values of raw data. P-values < 0.05 were considered significant. Survival plots were performed in GraphPad Prism version 7.0. Data for the indicated tumors was obtained from cBioportal (Cerami et al., 2012; Gao et al., 2013). Longitudinal and cross-sectional analysis of tumor volume where calculated using TumorGrowth tool using default parameters (Enot et al., 2018).

### Supplemental Tables

**Table S1:** Statistically significant terms upon GSEA analysis in melanoma versus nevi samples obtained from Kabbarah and Talantov data sets. Common terms are grouped; related to Figure 1 and Figure 2.

**Table S2:** Statistically significant terms upon GSEA analysis in shRRM2/shp16 versus shRRM2 alone (RNA-Seq); related to Figure 2.

**Table S3:** Statistically significant terms upon GSEA analysis in shRRM2/shp16 versus shRRM2 alone (RPPA); related to Figure 2.

**Table S4:** Relative abundance of nucleotides and deoxyribonucleotides; related to Figure 2.

**Table S5:** Leading-edge genes associated with “translation” term in shRRM2 versus shRRM2/shp16 GSEA used for survival analysis; related to Figure S2.

**Table S6:** Statistically significant terms upon GSEA analysis HRASV12/shRB versus HRASV12 alone in Chicas et al. data set; related to Figure S2.

**Table S7:** RT-qPCR expression analysis for the heavy and light polysome fractions; related to Figure 3.

**Table S8:** Isotopologue enrichment (M+5) after glucose labeling; related to Figure 3.

**Table S9:** Primers used for these studies.

**Figure S1.**
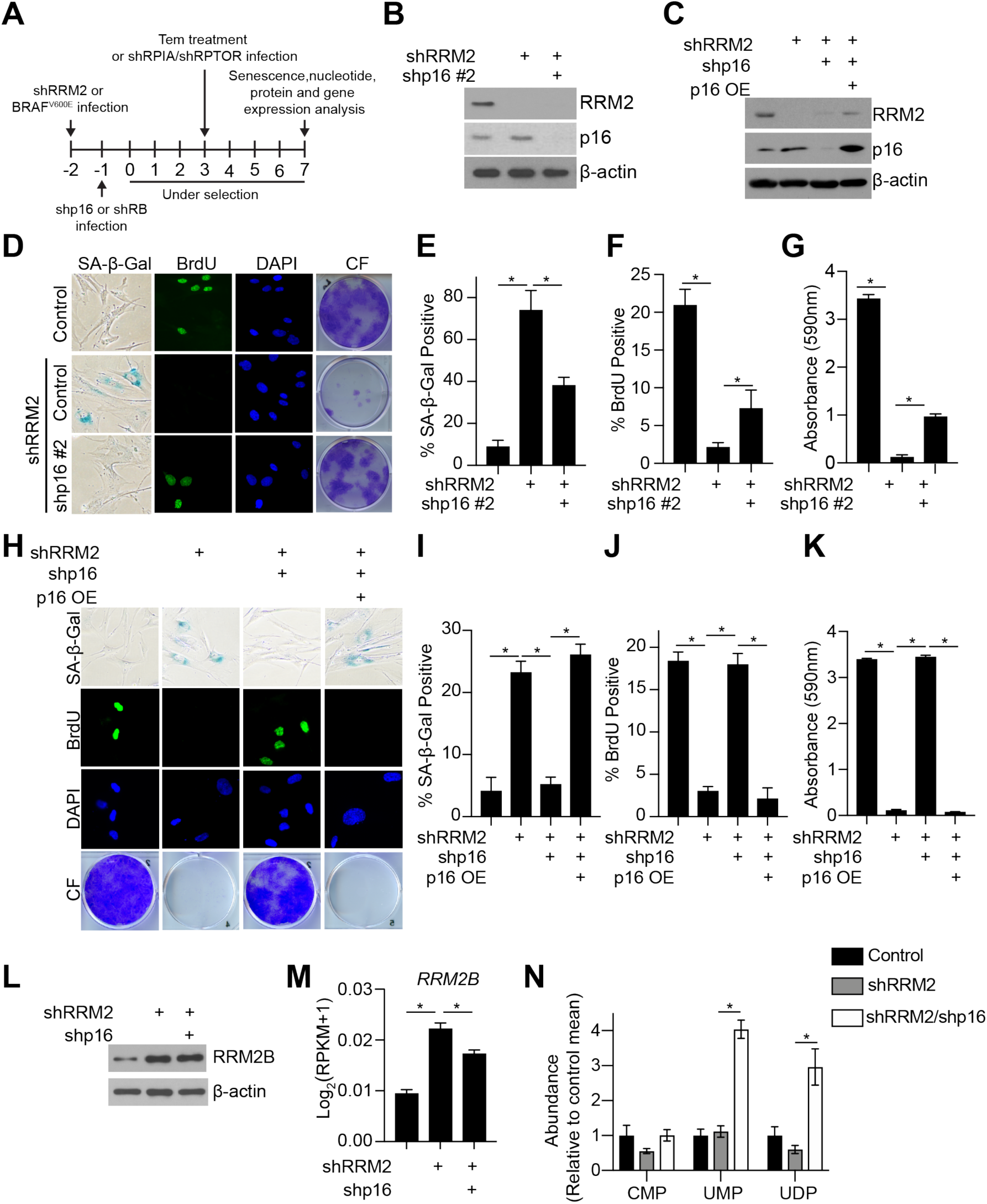
Overexpression of p16 represses senescence bypass; an independent shRNA targeting p16 also bypasses shRRM2-induced senescence; related to Figure 1. **(A)** Schematic of infection and treatment of cells. **(B-K)** Normal diploid IMR90 cells were infected with lentivirus expressing short hairpin RNAs (shRNAs) targeting RRM2 (shRRM2) alone or in combination with either an shRNA targeting p16 (shp16) or p16 overexpression cDNA construct. Empty vector was used as a control. **(B-C)** Immunoblot analysis of indicated proteins. One of 2 experiments is shown. **(D)** SA-β-Gal activity, BrdU incorporation, and colony formation. One of 2 experiments is shown. **(E)** Quantification of SA-β-Gal activity in (D). n=3/group, one of 2 experiments is shown. Data represent mean ± SEM. *p<0.001 **(F)** Quantification of BrdU incorporation in (D). n=3/group, one of 2 experiments is shown. Data represent mean ± SEM. *p<0.05 **(G)** Quantification of colony formation in (D). n=3/group, one of 3 experiments is shown. Data represent mean ± SEM. *p<0.001 **(H)** SA-β-Gal activity, BrdU incorporation, and colony formation. One of 2 experiments is shown. **(I)** Quantification of SA-β-Gal activity in (H). n=3/group, one of 2 experiments is shown. Data represent mean ± SEM. *p<0.001 **(J)** Quantification of BrdU incorporation in (H). n=3/group, one of 2 experiments is shown. Data represent mean ± SEM. *p<0.001 **(K)** Quantification of colony formation in (H). n=3/group, one of 3 experiments is shown. Data represent mean ± SEM. *p<0.001 **(L)** Immunoblot analysis of indicated proteins. One of 2 experiments is shown. **(M)** *RRM2B* expression from RNA-Seq analysis. *p<0.001 **(N)** Nucleotide analysis was performed by LC-HRMS in the indicated groups. n>3/group, one of at least 2 experiments is shown. Data represent mean ± SEM. *p<0.001

**Figure S2.**
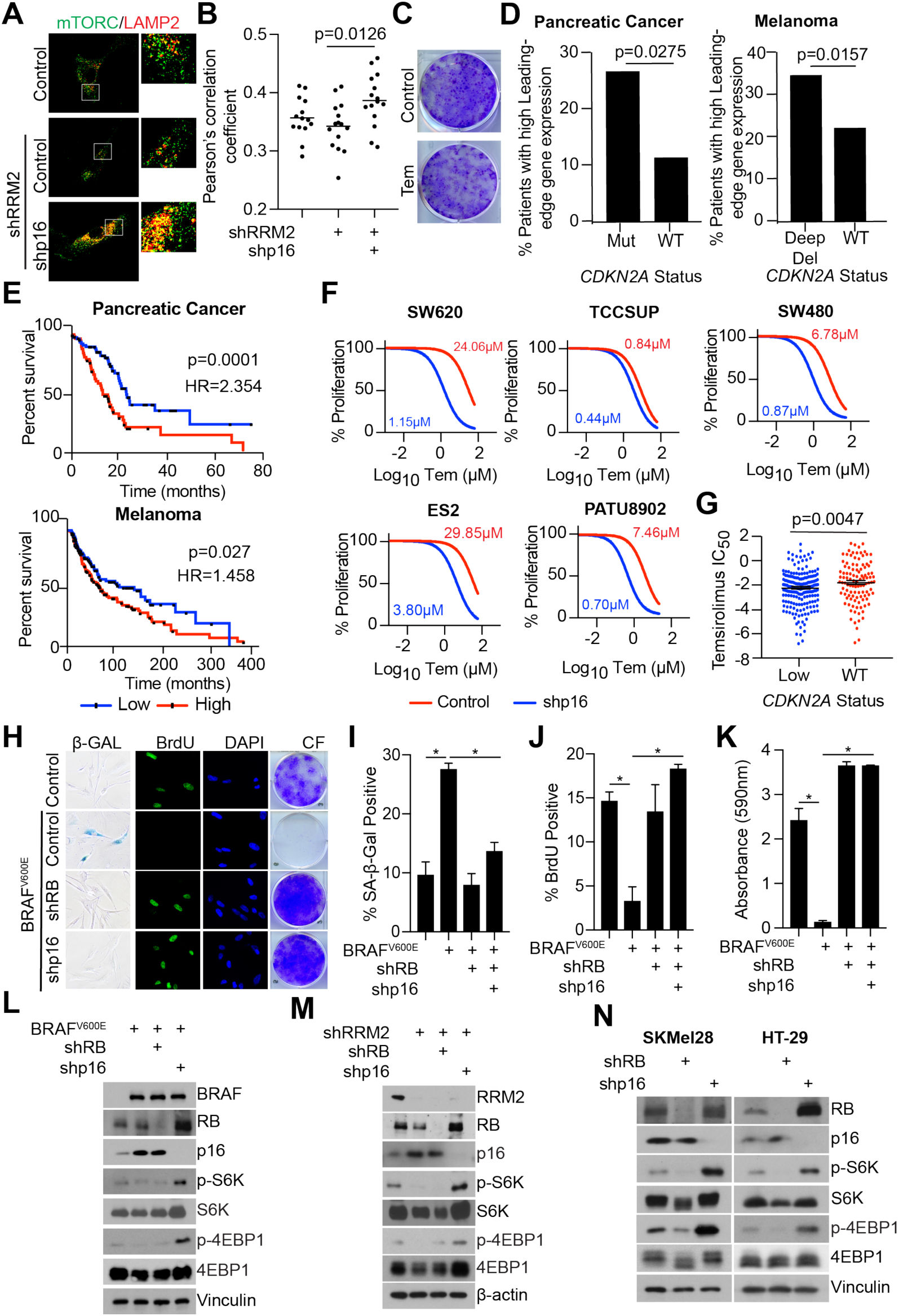
Suppression of p16 activates mTORC1 in a BRAF^V600E^ model of senescence bypass; suppression of RB does not activate mTORC1; related to Figure 2. **(A)** Normal diploid IMR90 cells were infected with lentivirus expressing short hairpin RNA (shRNAs) targeting RRM2 (shRRM2) alone or in combination with an shRNA targeting p16 (shp16). Empty vector was used as a control. Confocal microscopy images of co-localization of immunofluorescence staining using anti-mTOR and anti-LAMP2 antibodies. One of 2 experiments is shown **(B)** Quantification of (A). One of 2 experiments is shown. Data represent mean ± SEM. **(C)** Normal diploid IMR90 cells were treated with 0.5nM temsirolimus for 3 days and stained with 0.05% crystal violet. **(D)** Analysis of *CDKN2A* status of pancreatic cancer and melanoma TCGA data with high or low expression of leading-edge genes in “Translation” GSEA terms from RNA-Seq analysis (**Table S5**). **(E)** Kaplan Meier curves of overall survival for pancreatic cancer and melanoma patients with high or low expression of leading-edge genes in “Translation” GSEA terms from RNA-Seq analysis (**Table S5**). **(F)** The indicated cancer cell lines with wildtype p16 expression were infected with a short hairpin targeting p16 and then treated with a dose-course of temsirolimus under 0.5% FBS conditions. n=3/group, one of 2 experiments is shown. Data represent non-linear fit of transformed data. IC_50_ for each condition is indicated. **(G)** Analysis of *CDKN2A* copy number and temsirolimus IC_50_ data from the Dependency Map (depmap.org). **(H)** Normal diploid IMR90 cells were infected with retrovirus expressing BRAF^V600E^ alone or in combination with a lentivirus expressing shRNA targeting p16 (shp16) or RB (shRB). Empty vector was used as a control. SA-β-Gal activity, BrdU incorporation, and colony formation. One of 3 experiments is shown. **(I)** Quantification of SA-β-Gal activity in (H). n=3/group, one of 3 experiments is shown. Data represent mean ± SEM. *p<0.001 **(J)** Quantification of BrdU incorporation in (H). n=3/group, one of 3 experiments is shown. Data represent mean ± SEM. *p<0.003 **(K)** Quantification of colony formation in (H). n=3/group, one of 3 experiments is shown. Data represent mean ± SEM. *p<0.001 **(L)** Same as (H) but immunoblot analysis of the indicated proteins. One of 3 experiments is shown. **(M)** Normal diploid IMR90 cells were infected with lentivirus expressing shRRM2 alone or in combination with shp16 or shRB. Empty vector was used as a control. Immunoblot analysis of the indicated proteins. One of 3 experiments is shown. **(N)** p16 wildtype SKMel28 and HT-29 cancer cells were infected with lentivirus expressing shp16 or shRB. Immunoblot analysis of the indicated proteins. One of 3 experiments is shown.

**Figure S3.**
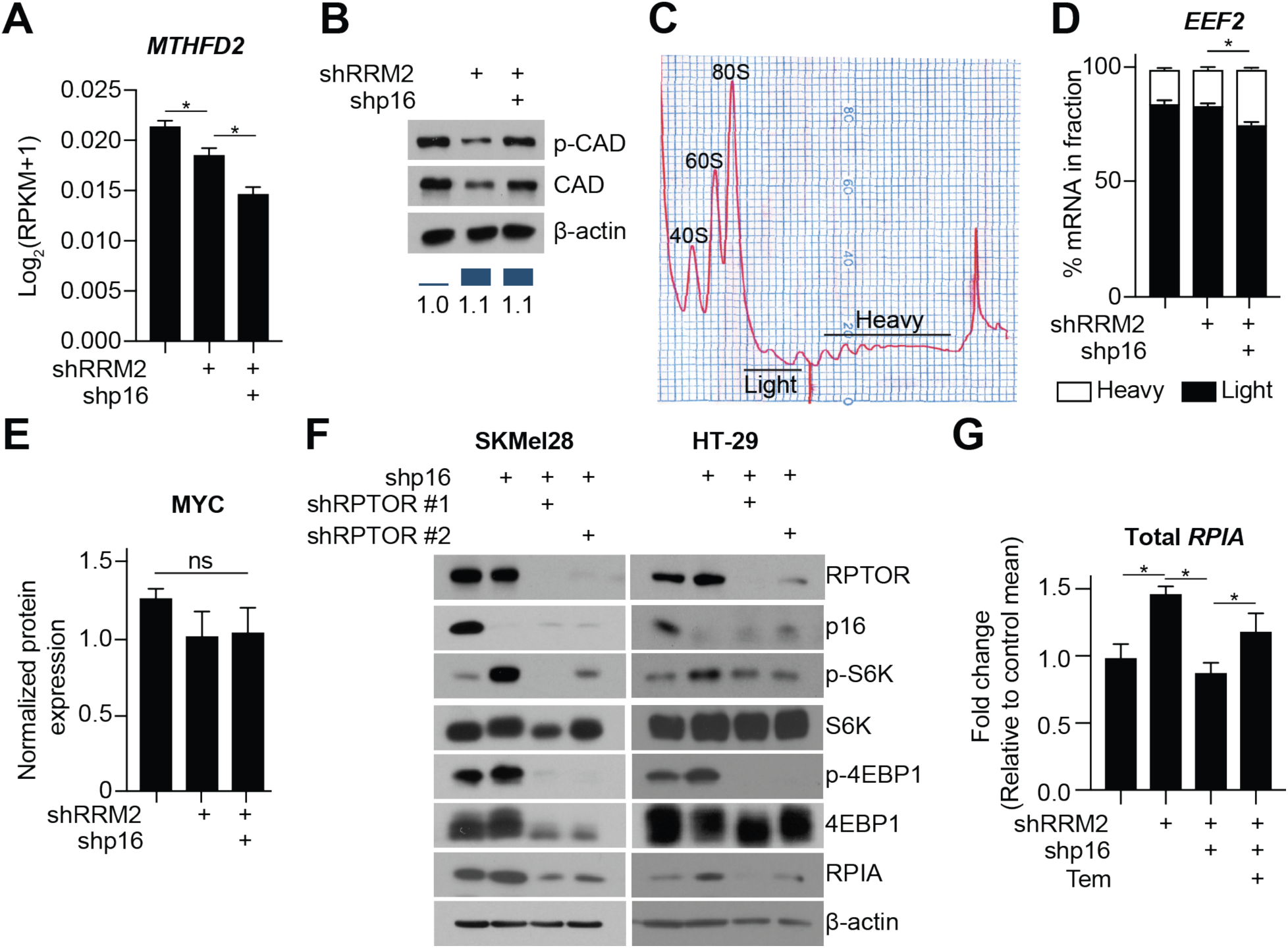
Suppression of p16 does not upregulate *MTHFD2* or CAD greater than shRRM2 controls; RAPTOR knockdown decreases RPIA expression; related to Figure 3. **(A-E)** Normal diploid IMR90 cells were infected with lentivirus expressing short hairpin RNA (shRNAs) targeting RRM2 (shRRM2) alone or in combination with an shRNA targeting p16 (shp16). Empty vector was used as a control. **(A)** *MTHFD2* expression from RNA-Seq analysis. n=3/group. Data represent mean ± SEM. *p<0.05 **(B)** Immunoblot analysis of p-CAD and total CAD. p-CAD fold change (FC) was performed relative to β-actin using ImageJ. **(C)** Representative polysome profile. “Light” and “Heavy” fractions are indicated. **(D)** Percentage of *EEF2* mRNA abundance in polysome fractions. n=3/group. Data represent mean ± SD. *p<0.001 **(E)** MYC protein expression in RPPA samples. ns=not significant **(F)** SKMel28 and HT-29 cells were infected with lentivirus expressing shp16 alone or in combination with two independent hairpins targeting RPTOR. Immunoblot analysis of the indicated proteins. One of 2 experiments is shown. **(G)** Same as (A) but cells were treated with Temsirolimus (Tem; 0.5nM) 4 days after starting selection. Total *RPIA* mRNA is shown. n=3/group, one of 2 experiments is shown. Data represent mean ± SD. *p<0.05

**Figure S4.**
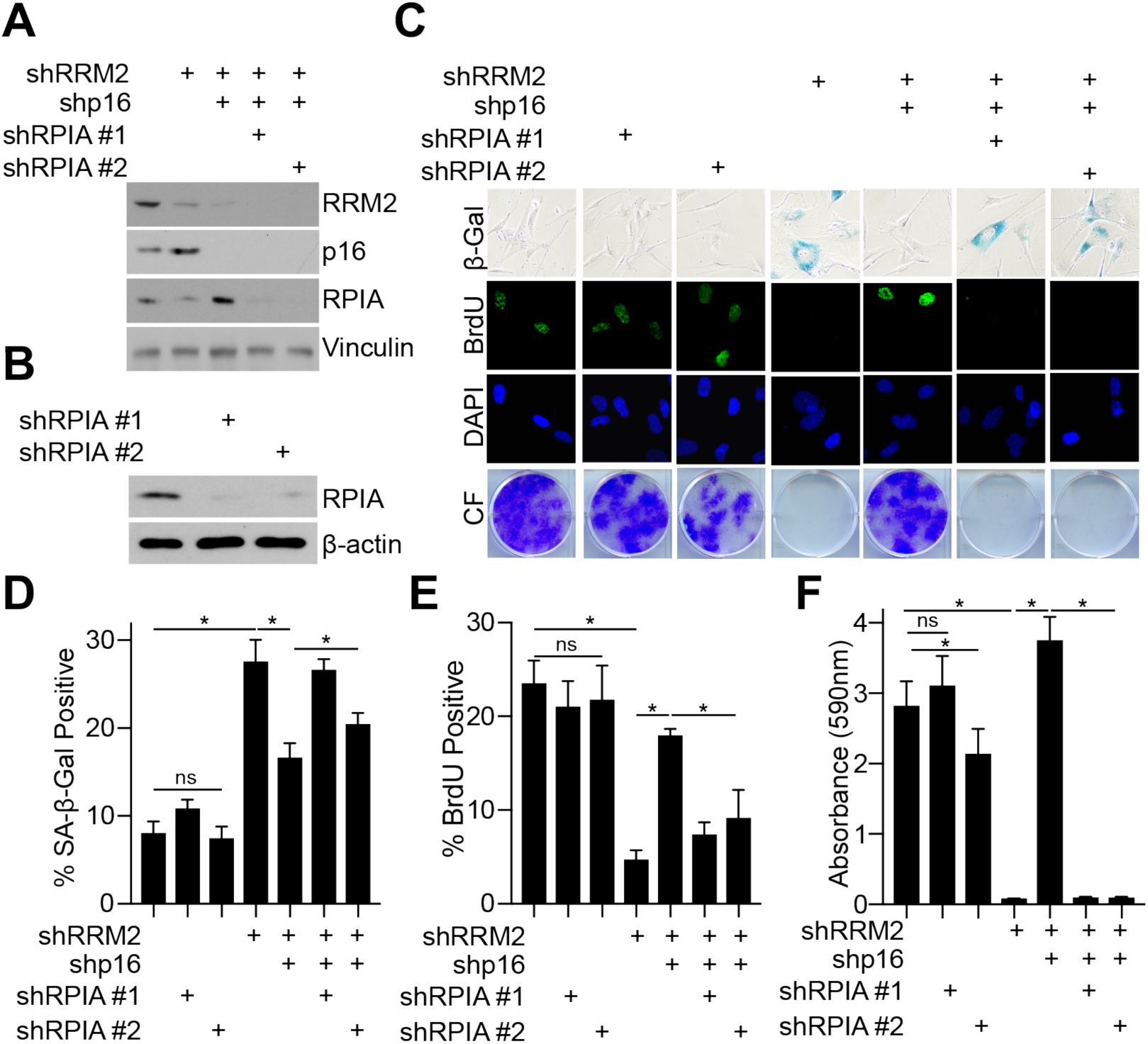
Knockdown of RPIA induces senescence only in shRRM2/shp16 and not parental IMR90 cells; related to Figure 4. **(A-F)** Parental or shRRM2/shp16 IMR90 cells were infected with lentivirus expressing 2 independent hairpins targeting RPIA. **(A-B)** Immunoblot analysis of the indicated proteins. One of 3 experiments is shown. **(C)** SA-β-Gal activity, BrdU incorporation, and colony formation. One of 3 experiments is shown. **(D)** Quantification of SA-β-Gal activity in (C). n=3/group, one of 3 experiments is shown. Data represent mean ± SEM. *p<0.001; ns=not significant **(E)** Quantification of BrdU incorporation in (C). n=3/group, one of 3 experiments is shown. Data represent mean ± SEM. *p<0.01; ns=not significant **(F)** Quantification of colony formation in (C). n=3/group, one of 3 experiments is shown. Data represent mean ± SEM. *p<0.05; ns=not significant

**Figure S5.**
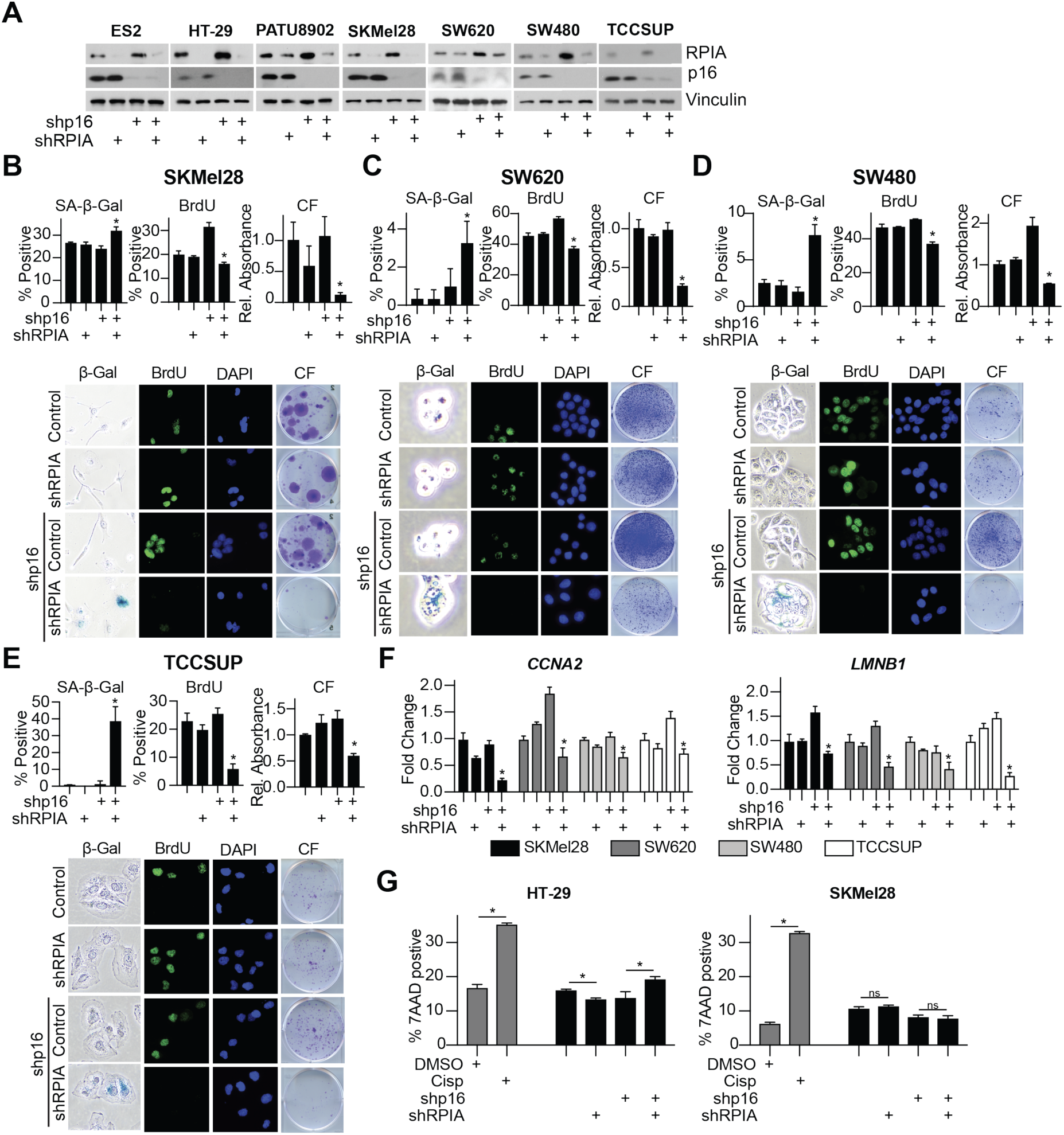
Knockdown of RPIA induces senescence only in shp16 cancer cells; related to Figure 4. **(A-C)** The indicated cancer cell lines with wildtype p16 expression were infected with a short hairpin targeting RPIA alone or in combination with a shRNA targeting p16. **(A)** RPIA and p16 western blot analysis of the indicated cell lines. Vinculin was used as a loading control. One of at least 2 experiments is shown. **(B-E)** SA-β-Gal activity, BrdU incorporation, and colony formation (CF) for each of the indicated cell lines. n=3/group, one of at least 2 experiments is shown. Data represent mean ± SEM. *p<0.05 vs. shp16 alone. **(F)** *CCNA2* and *LMNB1* expression was determined by RT-qPCR in the indicated cells. One of at least 2 experiments is shown. Data represent mean ± SD. *p<0.05 vs. shp16 alone. **(G)** SKMel28 and HT-29 cells were infected with a short hairpin targeting RPIA alone or in combination with a shRNA targeting p16 and analyzed for cell death using 7AAD staining followed by flow cytometry. Cisplatin was used as a positive control. *p<0.03; ns=not significant

